# Metabolomic Profiling Reveals Potential of Fatty Acids as Regulators of Stem-like Exhausted CD8 T Cells During Chronic Viral Infection

**DOI:** 10.1101/2024.10.07.617124

**Authors:** Katelynn R Kazane, Lara Labarta-Bajo, Dina R Zangwill, Kalle Liimatta, Fernando Vargas, Kelly C Weldon, Pieter C Dorrestein, Elina I Zúñiga

**Affiliations:** Division of Biological Sciences, University of California San Diego, La Jolla, California; Salk Institute for Biological Studies, La Jolla, California; Skaggs School of Pharmacy and Pharmaceutical Sciences, University of California San Diego, La Jolla, California

**Keywords:** Viral infection, metabolomics, fatty acids, CD8 T cell exhaustion, stem-like cells, LCMV

## Abstract

Chronic infections drive a CD8 T cell program termed T cell exhaustion, characterized by reduced effector functions. While cell-intrinsic mechanisms underlying CD8 T cell exhaustion have been extensively studied, the impact of the metabolic environment in which exhausted CD8 T cells (Tex) operate remains less clear. Using untargeted metabolomics and the murine lymphocytic choriomeningitis virus infection model we investigated systemic metabolite changes early and late following acute versus chronic viral infections. We identified distinct short-term and persistent metabolite shifts, with the most significant differences occurring transiently during the acute phase of the sustained infection. This included nutrient changes that were independent of viral loads and partially associated with CD8 T cell-induced anorexia and lipolysis. One remarkable observation was the elevation of medium- and long-chain fatty acid (FA) and acylcarnitines during the early phase after chronic infection. During this time, virus-specific CD8 T cells from chronically infected mice exhibited increased lipid accumulation and uptake compared to their counterparts from acute infection, particularly stem-like Tex (Tex^STEM^), a subset that generates effector-like Tex^INT^ which directly limit viral replication. Notably, only Tex^STEM^ increased oxidative metabolism and ATP production upon FA exposure. Consistently, short-term reintroduction of FA during late chronic infection exclusively improved Tex^STEM^ mitochondrial fitness, percentages and numbers. This treatment, however, also reduced Tex^INT^, resulting in compromised viral control. Our study offers a valuable resource for investigating the role of specific metabolites in regulating immune responses during acute and chronic viral infections and highlights the potential of long-chain FA to influence Tex^STEM^ and viral control during a protracted infection.

**Significance:** This study examines systemic metabolite changes during acute and chronic viral infections. Notably, we identified an early, transient nutrient shift in chronic infection, marked by an increase in medium- and long-chain fatty acid related species. Concomitantly, a virus-specific stem-like T cell population, essential for maintaining other T cells, displayed high lipid avidity and was capable of metabolizing exogenous fatty acids. Administering fatty acids late in chronic infection, when endogenous lipid levels had normalized, expanded this stem-like T cell population and enhanced their mitochondrial fitness. These findings highlight the potential role of fatty acids in regulating stem-like T cells in chronic settings and offer a valuable resource for studying other metabolic signatures in both acute and persistent infections.

## Introduction

Cytotoxic CD8 T cells are crucial players in antiviral immunity, essential for clearing acute viral infections and controlling persistent ones(1). However, long-term viral infections, such as Hepatitis B virus (HBV), Hepatitis C virus (HCV), and human immunodeficiency virus (HIV), can persist in the host by attenuating and evading the immune response(2). To cope with sustained viral loads, the body undergoes various adaptations, including attenuation of CD8 T cell functions, a phenomenon termed T cell exhaustion(3, 4). This adaptation was first identified in a mouse model of chronic lymphocytic choriomeningitis virus (LCMV) infection(5, 6) and has since been observed in other persistent infections, both in mice and humans(7–9), as well as in the tumor microenvironment(10, 11).

Exhausted CD8 T cells (Tex) are characterized by inhibited effector functions(5, 6), reduced proliferative capacity(12), impaired mitochondrial metabolism(13–16), and elevated expression of coinhibitory receptors such as PD-1, LAG3 and TIM3(3, 4). Additionally, Tex cells exhibit a unique transcriptional and epigenetic landscape and substantial heterogeneity(17–22). Early during viral infection, progenitor exhausted cells (Tex^STEM^) diverge from a short-lived effector-like cell population (Tex^EFF-LIKE^)(23, 24). With continuous antigen exposure, Tex^STEM^ differentiate into intermediate (Tex^INT^) and terminally exhausted (Tex^TERM^) populations(17, 18, 25–27). Tex^INT^ have a greater capacity for antiviral functions, producing cytokines and cytotoxic molecules that directly limit viral replication, while Tex^TERM^ are almost entirely dysfunctional(28, 29). Notably, Tex^STEM^ are present throughout infection, are essential for long-term immunity, and are the most responsive cell type to checkpoint immunotherapies(18, 27, 30). Furthermore, Tex^STEM^ exhibit the highest metabolic fitness among all Tex subsets(31). While the cell-intrinsic mechanisms driving T cell exhaustion have been extensively characterized(30, 32), the changes in the systemic milieu that exhausted CD8 T cells navigate—and how these changes impact different Tex subsets—remain less understood.

Metabolite concentrations are tightly regulated within the body, as they are essential drivers of fundamental cellular functions(33, 34). In addition to their clinical relevance as disease biomarkers, metabolites play a crucial role in regulating various biological processes, affecting transcriptional, proteomic, and epigenetic networks(33, 34). Interestingly, chronic infections have been shown to significantly impact the host metabolome(35–39). For example, individuals infected with HIV exhibit a sustained low tryptophan-to-kynurenine ratio and lipid dysregulation compared to HIV-negative individuals(37). Similar alterations in the systemic metabolic milieu have been reported in HBV-infected patients(36), in Plasmodium-infected humans and macaques(35), and in mice infected with Toxoplasma gondii(38) or LCMV(39).

In this study, we used untargeted metabolomics in the LCMV mouse model to unbiasedly compare side-by-side the systemic metabolome changes that take place early and late after acute vs. chronic viral infections. We found that the most significant metabolomic adaptations occurred during the early phase of chronic infection and were transient. These changes included shifts in nutrient availability that were uncoupled from viral loads and partly associated with CD8 T cell-dependent adipose tissue lipolysis and anorexia. A key observation was the temporary CD8 T cell-dependent elevation of medium- and long-chain fatty acid (FA) related species that was more pronounced early after chronic infection. Follow-up studies revealed that Tex cells exhibited greater lipid accumulation and uptake compared to their CD8 T cell counterparts in acute infection. Remarkably, the Tex^STEM^ subset showed enhanced lipid accumulation, long-chain FA uptake and metabolization compared to other Tex subsets. Consistently, reintroduction of FA at later stages of *in vivo* chronic infection improved mitochondrial fitness in only Tex^STEM^. This was accompanied with an exclusive increase in Tex^STEM^ percentages and numbers, while Tex^INT^ and Tex^TERM^ were reduced or unchanged, respectively. This Tex subset redistribution associated with increased viral burden upon FA administration. Our study provides a valuable resource to investigate the role of specific metabolites in antiviral immunity, which is exemplified by experiments underscoring the potential of long-chain FA as regulators of Tex subsets and viral control during chronic infection.

## Results

### Distinct systemic metabolic signatures characterized the early and late phases of acute and chronic LCMV infection

To define the metabolite composition of the milieu to which immune cells must adapt during acute and chronic viral infections, we investigated changes in plasma metabolites via untargeted metabolomics. We analyzed samples from mice infected with the persistent LCMV variant Clone 13 (Cl13) on day 8 (acute phase) and day 20 (chronic phase) post-infection (p.i.), and compared them to mice infected with the LCMV Armstrong 53b (ARM) variant that is cleared within a week(7), and to uninfected controls. At both time points, Principal Coordinate Analysis (PCoA) using Jaccard distances identified clusters based on infection type that were statistically distinct from each other and from uninfected mice (Fig. 1A, S1A&B). By day 8 p.i., LCMV ARM-infected mice exhibited changes in 256 molecular signatures compared to uninfected controls (Fig. 1B, Table S1). In contrast, LCMV Cl13 infection affected approximately 1.5 times more signatures, with 405 differential metabolomic changes observed (Fig. 1B, Table S2). By day 20 p.i., no differences were detected in ARM-infected vs. uninfected mice while 63 differential metabolomic signatures were observed in LCMV Cl13-infected vs. uninfected mice (Fig. 1B and Table S3). Of the metabolites perturbed in LCMV Cl13-infected mice, 356 (88%) were altered exclusively at day 8 p.i., while only 49 (12%) were persistently altered throughout day 20 p.i. (Fig. 1C, Table S4). Interestingly, this recovery from the initial chemical impact of LCMV Cl13 infection occurred despite sustained viral replication in the blood and most tissues(7, 40). This suggests that factors beyond high viral loads may drive the pronounced metabolic changes observed 8 days after chronic infection.

**Fig 1.**
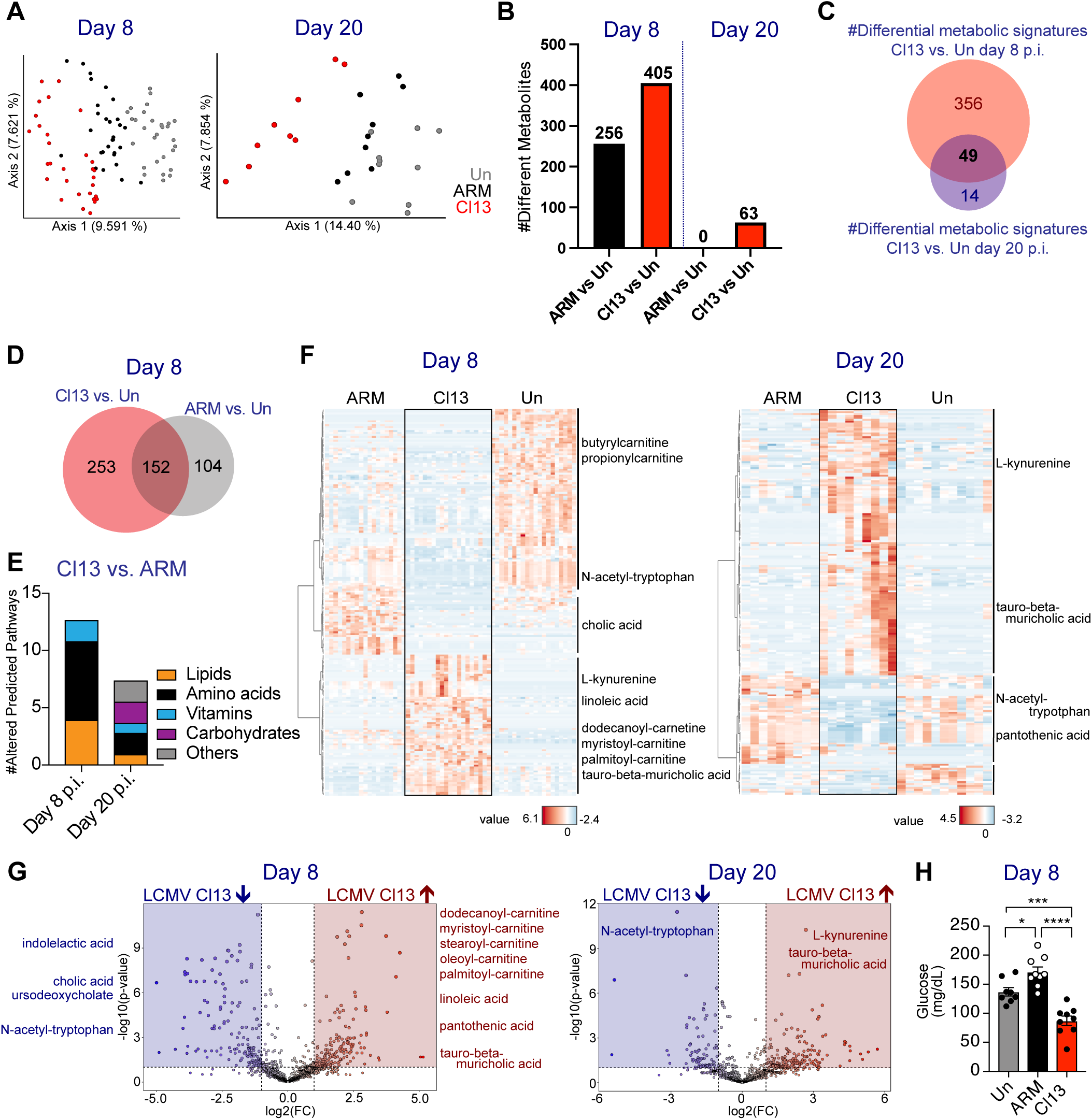
Distinct systemic metabolic signatures characterized the early and late phases of acute and chronic LCMV infection. C57BL/6 mice were infected with LCMV ARM, Cl13 or left uninfected (Un) and untargeted metabolomics was performed on plasma from days 8 and 20 p.i.. (A) PCoA plots with Jaccard distance on days 8 p.i. (left) and 20 p.i. (right). (B) Number of differential metabolites in ARM- or Cl13-infected mice vs. Un. (C) Overlap of differential metabolites in Cl13-infected vs. Un mice on days 8 and 20 p.i.. (D) Overlap of differential metabolites in Cl13-infected vs. Un and ARM-infected vs. Un on day 8 p.i.. (E) Functional activity prediction on metabolomes from Cl13-vs. ARM-infected mice. (F) Heatmap showing the top 200 differential metabolites ranked based on ANOVA f-value and MSI level 2 or 3 annotations for representative metabolites in plasma from Un, ARM- and Cl13-infected mice on day 8 p.i. (left) and day 20 p.i. (right). (G) Volcano plots showing the differential metabolites and MSI level 2 or 3 annotations for representative metabolites in plasma from ARM-vs. Cl13-infected mice on days 8 (left) and 20 p.i. (right). Blue and red boxes indicated downregulated and upregulated metabolites in LCMV Cl13 vs. ARM with a log2FC>2 and p-value<0.05. (H) Blood glucose levels in Un, ARM- and Cl13-infected mice on day 8 p.i.. (A-G) 2 pooled experiments with n=10-20 mice/group on day 8 p.i. and 1 experiment with n=9-10 mice/group on day 20 p.i.. (H) Representative of 2 experiments with n = 8-10 mice/group. (B-G) Wilcoxon rank test with false discovery rate (FDR)<0.05. (H) One-way ANOVA with Tukey’s correction. *p<0.05, ***<0.001, ****<0.0001.

To define specific metabolic differences occurring early and late after acute vs. chronic infection, we juxtaposed infection-specific chemical perturbations (defined in Fig. 1B, Table S1 and Table S2 by comparing infected vs. uninfected groups) and identified 253 metabolic signatures that were unique to LCMV Cl13-infected mice, 104 unique to LCMV ARM-infected mice and 152 overlapping between both groups of mice on day 8 p.i., which contain short-chain acyl carnitines and L-kynurenine, among other metabolites (Fig. 1D, Table S5). We also directly compared the metabolic signatures in LCMV ARM vs. Cl13 infected mice at day 8 (Table S6) and day 20 (Table S7) p.i. and identified 198 and 46 metabolites, respectively. We also performed Pathway Prediction Analysis(41) with the differential metabolic signatures in LCMV ARM-vs. Cl13-infected mice at day 8 (Table S6) and day 20 (Table S7) p.i., as determined by Wilcoxon rank test. This analysis revealed significant alterations in 13 pathways involving lipids, amino acids and vitamins at day 8 p.i., and 8 pathways at day 20 p.i. (Fig. 1E and Table S8). Subsequent Metabolomic Standard Initiative (MSI) level 2 and 3 annotations(42, 43) further confirmed that the metabolome differences after chronic vs. acute viral infection are time-point specific and involve heterogeneous groups of metabolites. This analysis identified metabolic biomarkers for LCMV ARM-infected mice(44) as well as LCMV Cl13-infected mice on days 8 and 20 p.i. (Fig. 1F, 1G and Tables S6 and S7, respectively). Notably, levels of many species related to lipid metabolism were changed at day 8 (but not day 20) following LCMV Cl13 compared to ARM infection. Changes included reductions in cholic acid levels, and increases in long-chain free FA (e.g. linoleic acid) and acylcarnitines (e.g dodecanoyl-carnitine, palmitoyl-carnitine and stearoyl-L-carnitine) (Fig. 1F&G, Tables S6&S7). These findings were consistent with the enhanced plasma FA elevation and adipose tissue lipolysis previously shown at days 6 through 8 following LCMV Cl13 infection(45), although a direct comparison to LCMV ARM-infected mice had not yet been made. We also detected a significant reduction in blood glucose (Fig. 1H) in day 8 (but not day 20) LCMV Cl13-infected compared to ARM-infected or uninfected mice (Tables S2-S7). Contrastingly, there were fewer metabolic signatures that remained altered from day 8 through day 20 post-LCMV Cl13 compared to ARM infection. Signatures that remained altered included elevated levels taurine-conjugated bile acids (tauro-beta-muricholic acid) as well as disrupted tryptophan metabolism, indicated by increased kynurenine and decreased N-acetyl-tryptophan (Fig. 1F&G, Tables S6&S7).

These data provide the first unbiased comprehensive characterization of the chemical milieu shaping the immune environment at different phases of chronic vs. acute viral infections. Our findings reveal that: i) the metabolic milieu differs significantly between chronic and acute infections, ii) the most pronounced chemical changes were uncoupled from pathogen load, occurring transiently during the acute phase of chronic infection, and iii) chronic infection induces a short-term shift in nutrient availability alongside sustained alterations in bile acids and tryptophan metabolism, among other metabolome disruptions.

### Transient elevation of medium- and long-chain fatty acid-related species was associated with CD8 T cell-induced adipose tissue lipolysis and anorexia after LCMV Cl13 infection

Given the increase in acylcarnitines and free FA early after LCMV Cl13 compared to ARM infection, we next investigated the kinetic of the systemic FA elevation and its potential association with sickness behaviors such as anorexia(45, 46) and associated adipose tissue lipolysis^41^. To do so, we first quantified plasma FA levels at various time points following both infections. We detected elevated FA concentrations in both LCMV ARM- and Cl13-infected compared to uninfected mice by day 7 p.i (Fig. 2A). However, LCMV Cl13 infection led to a greater and more sustained FA elevation through day 8 p.i., with levels returning to baseline by days 15 and 30 p.i. in both infection settings (Fig. 2A). This early plasma FA elevation was accompanied by a significant reduction in adipocyte area within the visceral white adipose tissue (WAT) (Fig. 2B), indicative of adipose tissue lipolysis, in LCMV Cl13-infected mice, which was consistent with previous results(45). We also detected adipocyte area reduction in LCMV ARM-infected mice by day 8 p.i., although the reduction was notably more pronounced in the chronic infection setting (Fig. 2B). In addition, the expression of lipase coding genes, *Pnpla2* (encoding ATGL) and *Lipe* (encoding HSL), were increased in the WAT from LCMV Cl13-infected mice (Fig. S2& (45)). WAT phosphorylation of HSL was also elevated in both infections but only reached significance in LCMV Cl13-infected mice by day 8 p.i. (Fig. 2C), further supporting increased lipolysis during chronic LCMV infection. Notably, the increase in systemic FA and reduction in adipocyte area were CD8 T cell-dependent, consistent with previous reports(45). CD8 T cell depletion resulted in a significant decrease in plasma FA levels (Fig. 2D& (45)) and increase in adipocyte area compared to mice with intact immune compartments (Fig. 2E& (45)). We further validated these findings by showing that HSL phosphorylation, but not total HSL, was reduced in CD8-depleted LCMV Cl13-infected mice (Fig. 2F). Overall, these results demonstrate that infection-induced elevation of FA is greater in persistent vs. acute LCMV infection, is associated with enhanced adipose tissue lipolysis in the chronic setting and is mediated by CD8 T cells.

**Fig 2.**
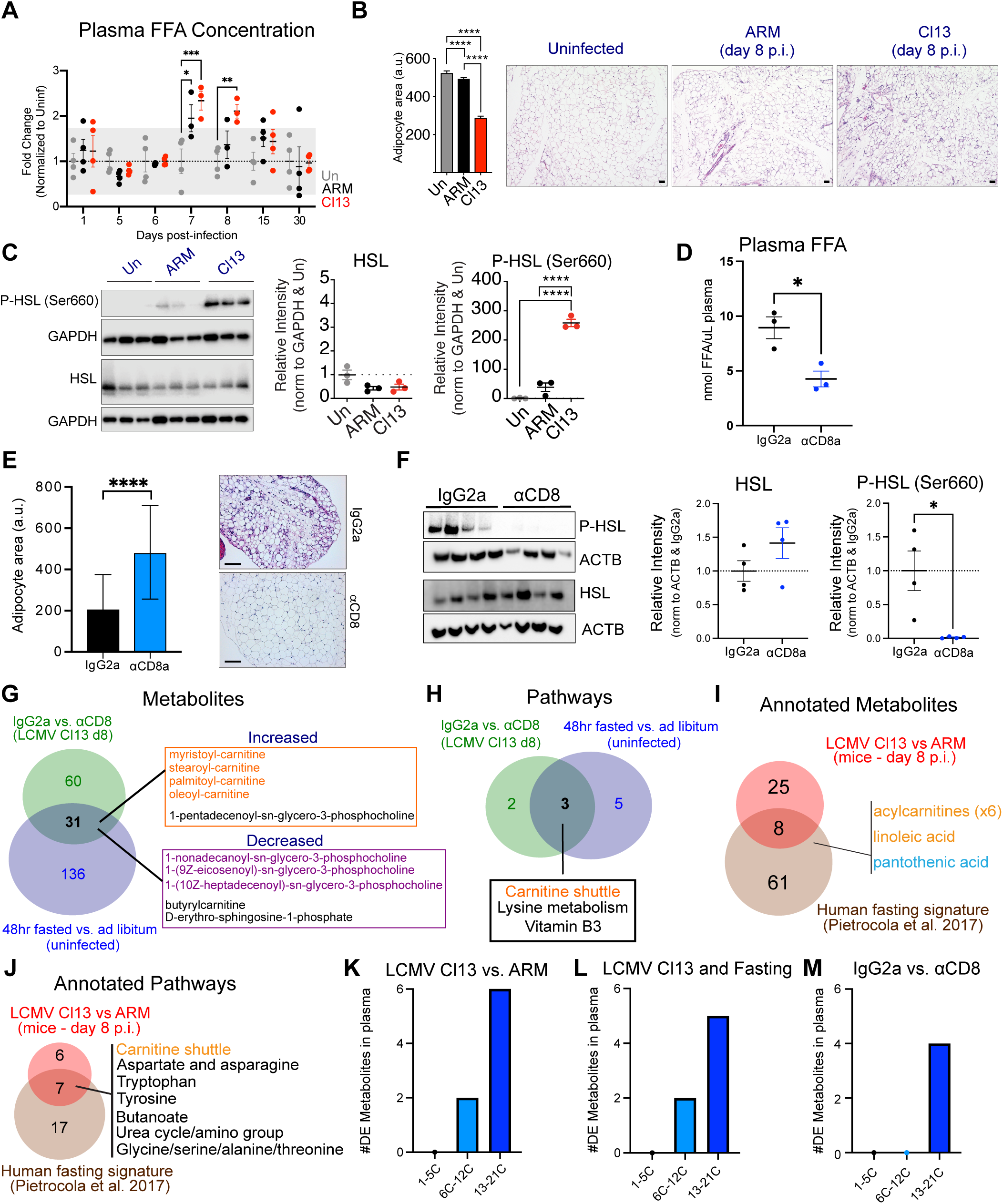
Elevation of long-chain FA related species was associated with CD8 T cell-induced adipose-tissue lipolysis and anorexia after LCMV Cl13 infection. C57BL/6 mice were infected with LCMV ARM, Cl13 or left uninfected (Un) and analyses were performed on days 8 and 20 p.i.. (A) Plasma free fatty acids (FFA) at indicated timepoints from days 1 through 30 p.i. normalized to Un levels. Horizontal lines indicate average Un (dotted line) ± 2 standard deviations (grey box). (B) Adipocyte area in visceral white adipose tissue (WAT) at day 8 p.i. and representative images (right). (C) Immunodetection of HSL and P-HSL (Ser660) in WAT at day 8 p.i.. Band intensity was normalized to GAPDH and Un mice. (D-H&M) Cl13-infected mice treated with isotype control (IgG2a) or anti-CD8-depleting antibodies (7CD8) and analyzed at day 8 p.i.. (D) Plasma FFA levels. (E) Adipocyte area in WAT and representative images (right). (F) Immunodetection of HSL and P-HSL (Ser660) in WAT. Band intensity was normalized to ACTB and isotype control mice. (G-H) Overlap of plasma metabolites (G) and predicted pathways (H) from isotype vs. anti-CD8 treated LCMV Cl13-infected mice and from 48hr fasted vs. ad libitum uninfected mice. (I-J) Overlap of plasma metabolites (I) and predicted pathways (J) from isotype vs. anti-CD8 treated LCMV Cl13-infected mice and human fasting metabolic signatures. (K – M) Number of differentially expressed metabolites with short-, medium-, or long-acyl chains that were elevated at day 8 p.i. in LCMV Cl13 vs ARM infected mice (K), LCMV Cl13-infected mice vs. human fasting signature (L), and isotype vs. anti-CD8 treated LCMV Cl13-infected mice (M). (A-F) Representative of 2 (B-F) or 3 (A) independent experiments with n=3-5 mice/group. (G-M) 3 pooled experiments with n=4-10 mice per group (CD8 depletion experiments) or 2 pooled experiments with n=9-10 mice/group (fasting experiments). (A-F) Averages ± SEM. (A) 2-way ANOVA. (B-C) One-way ANOVA with Tukey’s correction. (D-F) Two tailed Student’s t-test. *p<0.05, **<0.01, ***<0.001, ****<0.0001.

Previous studies, including our own, have described a profound and transient anorexia following LCMV Cl13 infection(45, 46), which we showed was dependent on CD8 T cells and absent in LCMV ARM-infected mice(46). Thus, to investigate the contribution of CD8 T cell induced anorexia to the systemic metabolic changes observed at day 8 after LCMV Cl13 infection, we evaluated CD8 T cell-dependent metabolic shifts during infection and compared them to metabolic signatures observed during fasting. Through untargeted metabolomics analysis of plasma from CD8-depleted vs. isotype-treated LCMV Cl13-infected mice, we identified 91 CD8 T cell-dependent metabolic signatures at day 8 p.i. (Table S9). These included CD8 T cell-dependent increases in acylcarnitines (i.e. palmitoyl-, oleoyl- and myristoyl-carnitine) and decreases in multiple phosphocholines (ie heptadecenoyl-, nonadecanoyl- and eicosenoyl-glycero-3-phosphocholine) (Fig. 2G, Table S9). We then compared these CD8 T cell-dependent metabolite shifts (Table S9) with those altered in uninfected mice fasted for 48hr (Table S10). We detected 31 overlapping metabolites (Table S11), including linoleic acid and several acyl-carnitines, such as palmitoyl- and stearoyl-carnitine, that were dependent on CD8 T cells after LCMV Cl13 infection and increased in fasted mice (Fig. 2G, Table S11). Pathway analysis further confirmed significant alterations in metabolites involved in the carnitine shuttle (Fig. 2H). Lastly, we compared the metabolic signatures and pathways uniquely altered in day 8 LCMV Cl13-infected mice (Tables S6&S8) vs. those altered in fasted human volunteers(47). This analysis revealed that 8 annotated (24.2%) metabolites and more than half of the pathways altered after LCMV Cl13 infection were similarly perturbed in both conditions across the two species, indicating conservation of metabolic changes induced by lack of food consumption (Fig. 2I&J, Table S12). Notably, consistent with the CD8-dependent metabolic shifts observed (Fig. 2G), several of the commonly altered metabolites were lipids, specifically within the acylcarnitine and free FA categories, including elevated levels of linoleic acid (Fig. 2I, Table S12). Moreover, as shown in Fig. 2J, the carnitine shuttle pathway also emerged as a major overlapping feature between LCMV Cl13-infected mice (Table S8) and fasted humans (Table S13). Overall, these results support CD8 T cell-induced anorexia as a contributor to the metabolome shifts occurring early after LCMV Cl13 infection, particularly influencing the elevation in systemic FA related species.

Finally, to better understand the specific lipids uniquely elevated in day 8 LCMV Cl13-infected mice, and their potential links to anorexia and CD8 T cell responses, we quantified the carbon number of each acyl group that showed significant alterations in LCMV Cl13 vs. ARM infection (Table S6) and overlapped with human fasting (Table S12) as well as CD8 T cell dependent metabolites (Table S9). Quantification of acyl group carbons in metabolites perturbed in day 8 LCMV Cl13 vs. ARM infection (Table S6) identified two medium-chain (6-12 carbons), and six long-chain (13-22 carbons) acyl-containing metabolites (Fig. 2K). Notably, the elevation in metabolites with medium- and long-chain acyl groups was common with human fasting (Fig. 2L), while the increase in metabolites with long-chain acyl groups was mostly CD8 T cell-dependent (Fig. 2M).

Collectively, these data suggest a model in which CD8 T cell-induced adipose tissue lipolysis(45) and anorexia(45, 46) early in LCMV Cl13 infection partly drive the transient plasma metabolite changes observed. This includes elevation of medium and long-chain FA and acyl-carnitines, drawing some parallels to human fasting conditions. Our results also suggest that adipose tissue lipolysis may be triggered after acute infection, though to a much lesser extent, likely due to the absence of anorexia in this context(46).

### Tex^STEM^ exhibited enhanced lipid content and uptake after LCMV Cl13 infection

FA can serve as a fuel source for CD8 T cell metabolism(48). After observing the significant systemic elevation of medium- and long-chain FA and derived acylcarnitines at day 8 following LCMV Cl13 vs. ARM infections (Fig. 2K, Table S6), we investigated their potential accumulation and uptake by virus-specific CD8 T cells, which peak around this time point(5). Notably, virus-specific (D^b^/GP_33-41_ tetramer-positive) CD8 T cells from LCMV Cl13-infected mice displayed higher neutral lipid content (Fig. 3A) and greater uptake capacity for both medium- and long-chain FA (Fig. 3B&C) compared to LCMV ARM-infected mice. To determine which CD8 T cell subpopulation exhibited more profound lipid-avid phenotype following LCMV Cl13 infection, we analyzed the lipid content and uptake capacity of splenic Tex subsets at day 8 p.i.. Tex populations were categorized into Tex^STEM^ and Tex^EFF-LIKE^, distinguished by the expression of *Slamf6* (LY108). Additionally, Tex^EFF-LIKE^ were further subdivided based on CX3CR1 expression, which differentiates more functional Tex cells (Tex^INT^) later in infection(24, 28, 29). Among the splenic Tex subsets, Tex^STEM^ exhibited the highest neutral lipid content at day 8 p.i. (Fig. 3D). Notably, Tex^STEM^ also showed the greatest capacity for medium- and long-chain FA uptake among all Tex subsets analyzed (Fig. 3E&F). Overall, these findings showed that, early in a persistent infection when endogenous FA are present at elevated levels in circulation, virus-specific CD8 T cells, particularly the Tex^STEM^ population, displayed increased lipid content as well as FA uptake capacity, raising the possibility that FA exposure might be able to fuel Tex^STEM^ metabolism.

**Fig 3.**
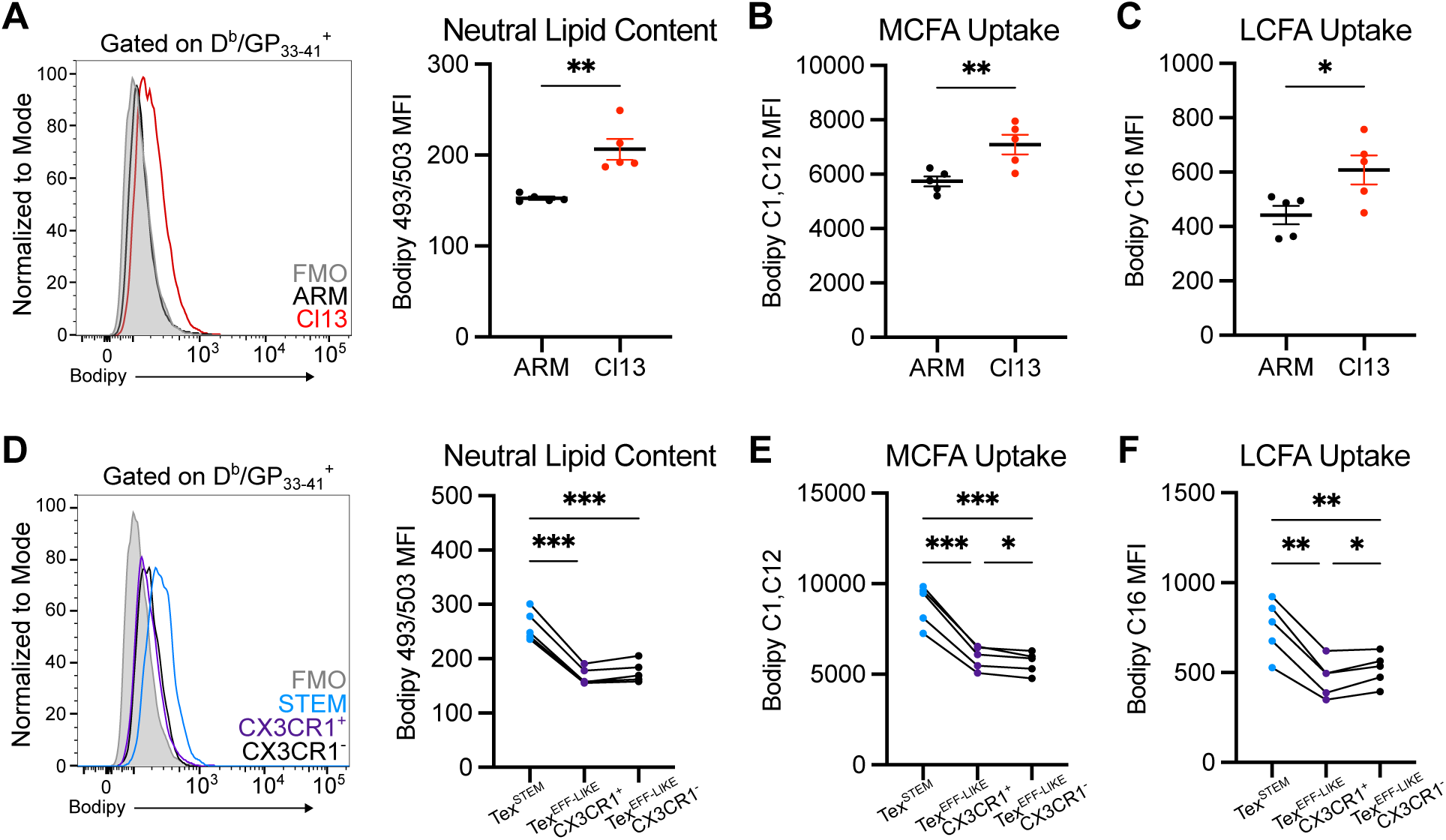
Tex^STEM^ showed higher neutral lipid content and FA uptake than other Tex subsets after LCMV Cl13 infection. C57BL/6 mice were infected with LCMV ARM or Cl13 and splenocytes were isolated and stained with BODIPY 493/503, BODIPY C1,C12, or BODIPY C16 at day 8 p.i.. Neutral lipid content (A,D), medium-chain FA uptake (B,E) and long-chain FA uptake (C,F) in CD8^+^ D^b^/GP_33-41_ tetramer^+^ cells from LCMV ARM vs. Cl13 infected mice (A-C) and across CD8^+^ D^b^/GP_33-41_ tetramer^+^ Tex^STEM^ (LY108^+^ CX3CR1^-^) and Tex^EFF^ (LY108^-^ CX3CR1^+^ or LY108^-^ CX3CR1^-^) from LCMV Cl13-infected mice. (A-C) Averages ± SEM. (D-F) Values from different Tex subsets corresponding to the same mouse are connected with a line. (A-F) Data are representative of two (A-C) or three (D-F) experiments, with n=4-5 mice/group. (A-C) Two-tailed Student’s t-test. (D-F) One-way ANOVA with Tukey’s correction. *p<0.05, **<0.01, ***<0.001.

### *Ex vivo* FA exposure induced gene expression changes related to lipid metabolism across all Tex subsets

As Tex populations, particularly Tex^STEM^, were more avid for medium- and long-chain FA uptake than their counterparts after acute LCMV infection (Fig. 3), we investigated the direct transcriptional impact of FA exposure on each Tex subset using RNA sequencing (RNAseq). We incubated FACS-purified Tex^STEM^ and Tex^EFF-LIKE^ subsets isolated from the spleens of LCMV Cl13-infected mice at day 8 p.i. (Fig. S3A) with the 16-carbon long-chain FA palmitate, which corresponding acylcarnitine (palmitoylcarnitine) is increased in a CD8 T cell-dependent manner following LCMV Cl13 infection (Tables S2, S6&S9). Palmitate was conjugated with bovine serum albumin (BSA) to mimic how cells encounter circulating FA, which are typically transported by albumin(49). Cells were incubated with either palmitate-BSA or BSA alone for 8 hours before RNA isolation and subsequent RNAseq. Principle component analysis (PCA) revealed that the Tex^STEM^ and Tex^EFF-LIKE^ populations clustered separately (Fig. 4A), suggesting that FA exposure did not override their original transcriptional identity. Tex^STEM^ cells exhibited differential expression of 43 genes upon FA incubation (Fig. 4B, Table S14), including up-regulation of *Cpt1a,* which encodes carnitine palmitoyltransferase 1A (CPT1A), and *Slc25a20*, which encodes carnitine-acylcarnitine translocase (CACT)(50). These proteins are crucial components of the carnitine shuttle, facilitating transport of FA into the mitochondria where they undergo FAO(51, 52). On the other hand, Tex^EFF-LIKE^ CX3CR1^+^ and CX3CR1^-/low^ displayed a total of 135 and 78 significantly perturbed genes, respectively (Fig. 4B, Table S14).

**Fig 4.**
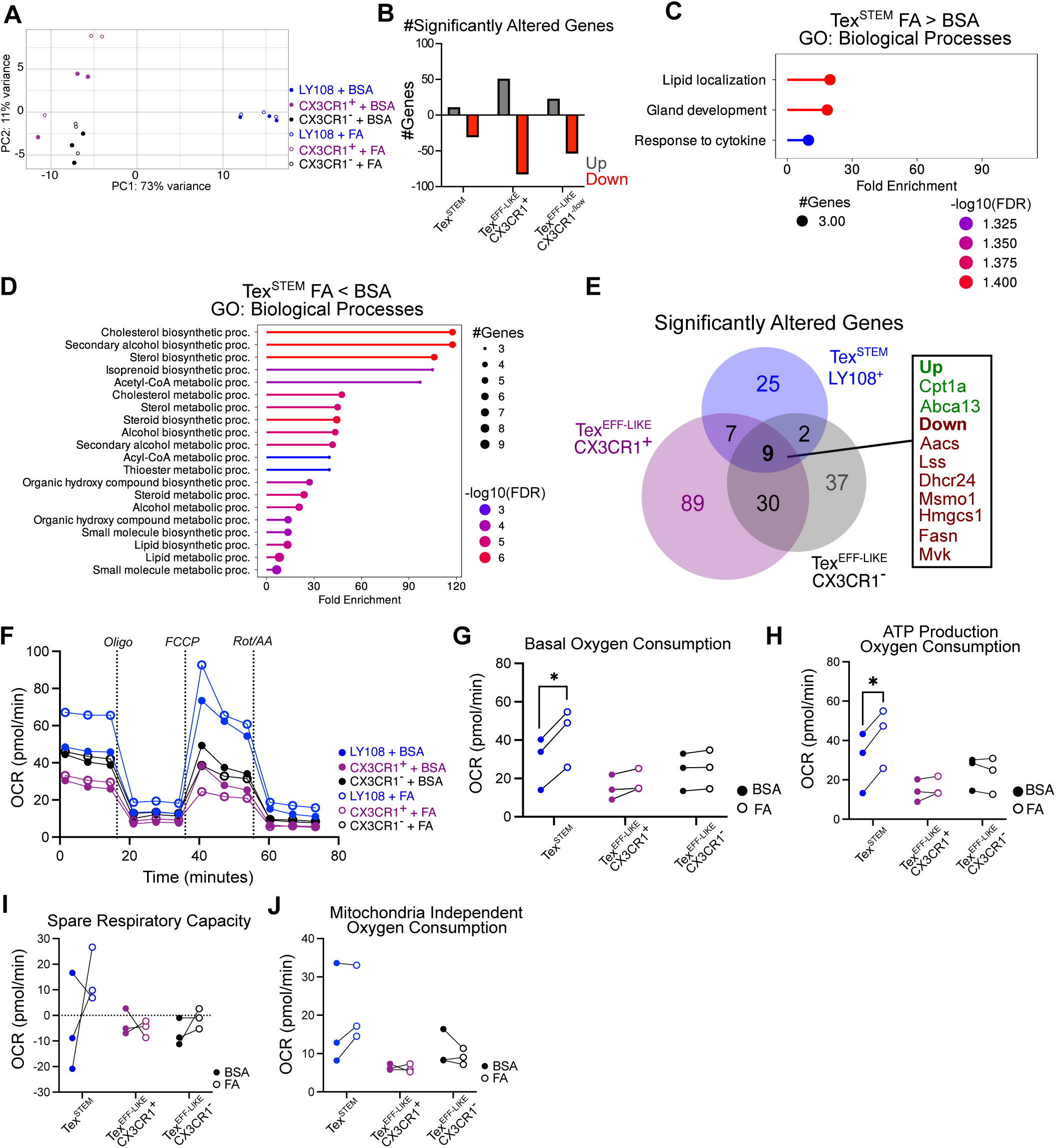
*Ex vivo* FA exposure altered lipid metabolism related transcripts across Tex subsets and increased oxygen consumption in Tex^STEM^ from LCMV Cl13 infected mice. C57BL/6 mice were infected with LCMV Cl13, and CD8^+^ PD1^+^ Tex^STEM^ (LY108^+^ CX3CR1^-^) or Tex^EFF-LIKE^ (LY108^-^ CX3CR1^+^ or LY108^-^ CX3CR1^-/low^) splenocytes from day 8 p.i. were FACS-purified and incubated with palmitate-BSA (FA) or BSA alone (BSA). (A-E) Eight hours post-culture, Tex subsets were processed for RNAseq. (A) PCA plot showing individual Tex subsets after BSA vs. FA incubation. (B) Number of differentially expressed genes (p-value<0.05, log2FC>0.5) in BSA vs. FA incubated Tex subsets. (C-D) GO biological processes enriched by upregulated (C) or downregulated (D) genes after FA vs. BSA incubation. (E) Venn diagram showing overlap of differentially expressed genes in BSA vs. FA incubated in different Tex subsets. (F) Seahorse assay traces of oxygen consumption rates (OCR) over time before and after the addition of Oligomycin (Oligo), FCCP and Rotenone/Antimycin A (Rot/AA). (G-J) Average basal oxygen consumption rate (G), ATP-production linked oxygen consumption (H), spare respiratory capacity (I), and mitochondrial-independent oxygen consumption (J) in each Tex subset incubated with BSA vs. FA. (A-E) Data pooled across 3 independent repeats with n=4-5 mice/experiment. (F) Data representative of 3 individual repeats with n=8-10 mice/experiment. (G-J) Each data point represents a single experimental repeat, paired data from the same repeat are connected with a line. (G-J) Paired student’s t-test. *p< 0.05, **<0.01, ***<0.001.

Differentially expressed genes (DEGs) from individual Tex subsets were next submitted for pathway analysis using gene ontology (GO) and the Kyoto Encyclopedia of Genes and Genomes (KEGG) with a false discovery rate cutoff of 0.05. Only enriched GO terms with more than 3 DEG were included. FA treatment in Tex^STEM^ (Fig. 4C) and Tex^EFF-LIKE^ CX3CR1^-/low^ (Fig. S3B) upregulated expression of genes related to lipid localization, while Tex^EFF-LIKE^ CX3CR1^+^ cells were enriched for the KEGG pathway related to Peroxisome Proliferator-Activated Receptor (PPAR) signaling (Fig. S3C), which is a master regulator of cellular metabolism, including FAO(53). On the other hand, all Tex subsets demonstrated consistent downregulated expression of genes encoding cholesterol biosynthetic processes and steroid metabolic processes (Fig. 4D, Fig. S3B & C).

To better understand shared FA effects in all Tex subsets, we then compared the genes significantly altered by the addition of palmitate (adjusted p-value < 0.05) and identified 9 DEGs that were consistently altered in the same direction in all three Tex subsets (Fig. 4E). Among these, *Cpt1a* and *Abca13* were upregulated. *Abca13* encodes for ATP Binding Cassette Subfamily A Member 13 (ABCA13) which functions as a lipid shuttle, mediating lipid transport across cell membranes(54). Conversely, the downregulated genes included those involved in FA synthesis, such as *Fasn*(*55*), and cholesterol biosynthesis, including *Aacs, Lss, Dhcr24, Msmo1, Hmgcs1* and *Mvk*(*56*). Thus, these results demonstrate that FA consistently alter the expression of lipid metabolism-related genes in all Tex subsets, suggesting that FA exposure during infection could directly modulate the transcriptional profile of exhausted CD8 T cells.

### *Ex vivo* FA exposure enhanced Tex^STEM^ oxidative metabolism and ATP production

Given the increased availability of long-chain acyl group-containing metabolites in the plasma of day 8 LCMV Cl13-infected mice and the impact of palmitate on Tex transcriptional profiles, particularly the shared upregulation of *Cpt1a*, we assessed the capacity of Tex subsets to metabolize long-chain FA. We provided FACS-purified Tex subsets, isolated at day 8 post-LCMV Cl13 infection (Fig. S3A), with palmitate and used the Agilent Seahorse platform to measure the oxygen consumption rate (OCR) as a proxy for oxidative phosphorylation(57). Consistent with previous reports(31), Tex^STEM^ demonstrated higher OCR compared to Tex^EFF-LIKE^ subsets (Fig. 4F&G). Notably, we observed that among the three Tex subsets analyzed, only the Tex^STEM^ subset exhibited an increase in basal OCR upon incubation with palmitate (Fig. 4F&G). This was accompanied by an increase in ATP production, as indicated by the decrease in OCR following the inhibition of ATP synthase activity with Oligomycin (Fig. 4F&H). In contrast, spare respiratory capacity (SRC), defined as the difference between basal OCR and maximal OCR after mitochondrial uncoupling with FCCP, remained unaffected by palmitate addition across all subsets. Similarly, non-mitochondrial oxygen consumption, indicated by the OCR changes after the addition of Rotenone and Antimycin A (Rot/AA), which inhibit complexes I and III of the electron transport chain, was also unchanged (Fig. 4I&J). These results indicate that Tex^STEM^ was the only Tex subset able to effectively metabolize long-chain FA from the environment, which may be partly due to the aforementioned enhanced capacity of Tex^STEM^ cells to uptake medium- and long-chain FA.

### *In vivo* FA administration enhanced Tex^STEM^ mitochondrial fitness during LCMV Cl13 infection

After demonstrating the superior ability of Tex^STEM^ to uptake and metabolize FA *ex vivo*, we investigated the effect of *in vivo* FA administration on shaping Tex subsets during persistent viral infection. To this end, we intraperitoneally injected LCMV Cl13-infected mice with a 1:1 mixture of lauric and palmitic acids diluted in castor oil from days 16-20 p.i. (Fig. 5A), a period when endogenous FA are no longer elevated (Fig. 2A). At this timepoint, the Tex^STEM^ population remains present and is crucial for sustaining a pool of effector-like Tex^INT^ cells(30). To have an overall picture of the effect that the *in vivo* FA inoculation may have on each Tex subset, we first performed RNAseq in Tex subpopulations FACS-purified one day after completion of the treatment (Fig. S4A). PCA indicated that, as in the *ex vivo* treatment setting, Tex subsets clustered independently, regardless of FA treatment, indicating that FA did not alter the Tex subset identity (Fig. 5B). We then compared the perturbed genes across Tex subsets. Tex^STEM^ was greatly altered by the FA administration, with 159 upregulated and 136 downregulated genes (Fig. 5C, Table S15). Upregulated genes were associated with GO biological processes related to immune cell apoptosis, regulation of gene expression and protein metabolic processes (Fig. 5D). Downregulated genes included those related to DNA replication and cell division, as well as cell cycle related genes (Fig. 5E). Although there were 27 shared genes across all three subsets (Fig. 5F, Table S16), there were no upregulated GO terms or KEGG pathways associated with these genes. However, GO biological processes related to DNA replication and cell cycle processes were both downregulated in the shared DEG list (Fig. S4B).

**Fig 5.**
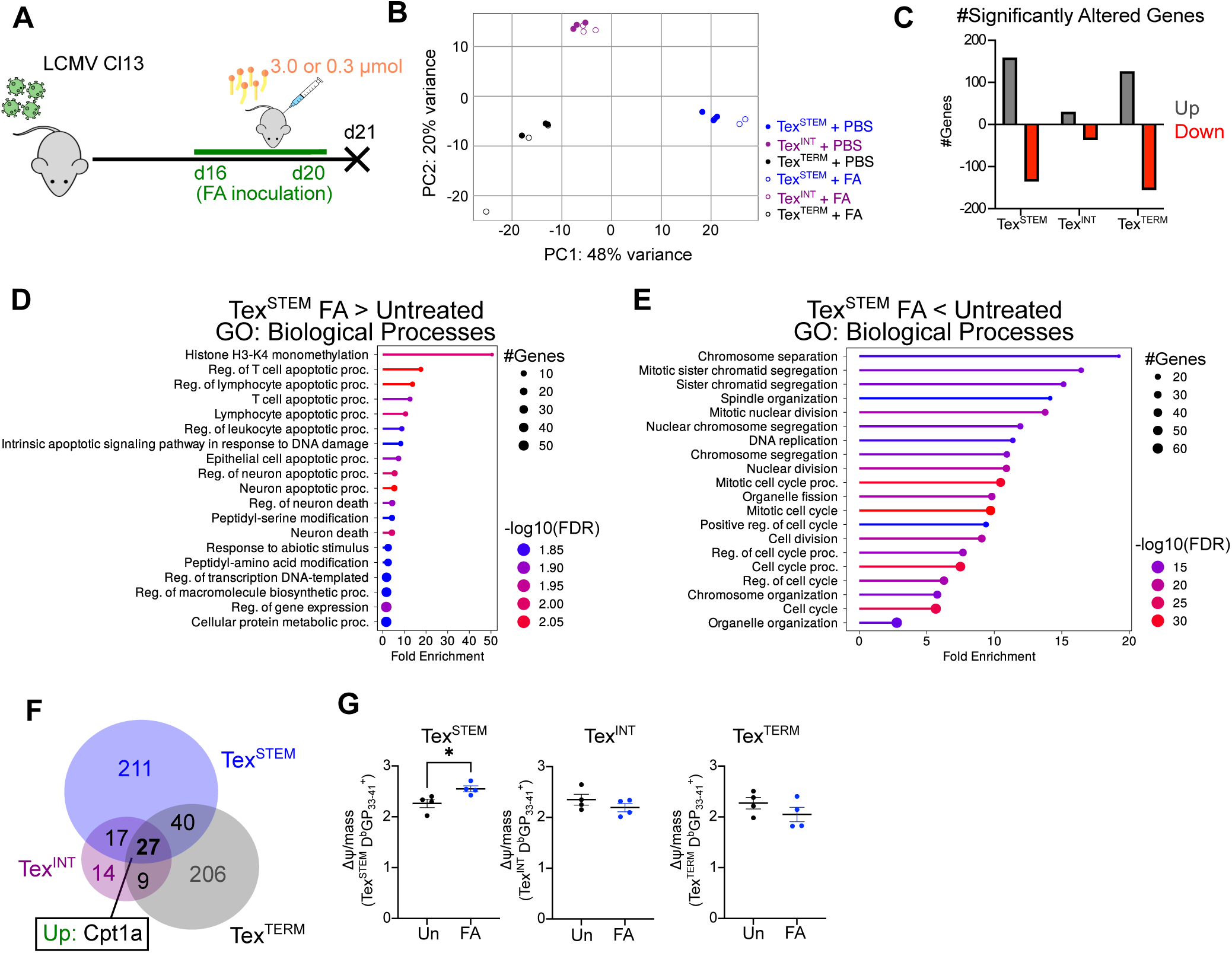
*In vivo* FA administration enhanced *Cpt1a* gene expression across Tex subsets and enhanced Tex^STEM^ mitochondrial fitness during LCMV Cl13 infection. C57BL/6 mice were infected with LCMV Cl13 and treated i.p. with 3.0 μmol or 0.3 mmol of FA (1:1 lauric and palmitic acid) or PBS every 12 hours from days 16 to 20 p.i.. CD8^+^ PD1^+^ Tex^STEM^ (LY108^+^ CX3CR1^-^), Tex^INT^ (LY108^-^ CX3CR1^+^) or Tex^TERM^ (LY108^-^ CX3CR1^-^) were FACS purified for RNAseq (B-F) or stained for FACS analysis (G) at day 21 p.i.. (A) Experimental design. (B) PCA showing individual Tex subsets from FA-treated vs. untreated mice. (C) Number of differentially expressed genes (p-value<0.05, log2FC>0.5). (D,E) GO biological processes enriched by upregulated (D) or downregulated (E) genes in Tex^STEM^ from FA treated vs. untreated mice. (F) Venn diagram showing overlap of differentially expressed genes in Tex subsets from FA treated vs. untreated mice. (G) Averages ± SEM of mitochondrial polarization (charge/mass) in CD8^+^ D^b^/GP_33-41_ tetramer^+^ Tex subsets. (A-F) Data are combined across three experiments with n=4-5 mice/group. (G) Data are representative of three experiments with n=4-5 mice/group. Unpaired two-tailed Student’s t-test. *p<0.05.

Notably, consistent with *ex vivo* gene induction (Fig. 4E), all Tex subsets exhibited a shared upregulation of *Cpt1a* upon *in vivo* FA treatment (Fig. 5F, Table S16), suggesting increased transcription of components of the carnitine shuttle and improved FA transportation into the mitochondrion for FAO(51). Given this observation, we next evaluated the mitochondrial polarization normalized to the mitochondrial mass in the three Tex subsets after *in vivo* FA administration(58). This parameter reflects the proton gradient across the mitochondrial inner membrane, which can be used as a proxy for mitochondrial activity(59). We detected enhanced mitochondrial polarization in virus-specific Tex^STEM^, but not in other Tex subsets, upon FA inoculation (Fig. 5G), suggesting that Tex^STEM^ may be better equipped than other Tex subsets to process the systemic FA provided *in vivo*.

Together with the enhanced FA uptake capacity (Fig. 3E&F) and FA metabolization (Fig. 4G&H), these findings are consistent with a model in which the Tex^STEM^ robust capacity for FA uptake and oxidation position them as the primary beneficiaries of systemic FA availability, which selectively enhances their mitochondrial fitness.

### *In vivo* FA administration increased Tex^STEM^ while reducing Tex^INT^ and impairing viral control during LCMV Cl13 infection

Tex^STEM^ are a self-renewing population, and their differentiation is crucial for sustaining a pool of effector-like Tex^INT^ cells throughout persistent viral infection(30). Given the FA improved mitochondrial activity in Tex^STEM^ (Fig. 5G), we next evaluated the impact of the *in vivo* FA treatment on the proportion and number of each Tex subset using two different doses (i.e. 0.3 μmol & 3 μmol) of a 1:1 mixture of lauric and palmitic acids diluted in castor oil and inoculated intraperitoneally. One day after the treatment concluded, we detected a significant increase in the percentage and number of virus-specific Tex^STEM^ cells (Fig. 6A&B), which was consistent with their improved mitochondrial activity (Fig. 5G). The FA treatment, however, also decreased the proportion and numbers of Tex^INT^ cells (Fig. 6A&C), while Tex^TERM^ cells were not significantly altered (Fig. 6A&D). We did not detect any difference in interferon-γ production upon *ex vivo* peptide stimulation (Fig. S5A) or TIM-3 expression within the different Tex subsets (Fig. S5B). There was, however, a significant increase in PD-1 across all Tex subsets (Fig. 6E), which could be associated with potentially higher viral loads(60). Indeed, quantification of viremia the day after the FA treatment was completed demonstrated increased viral titers in the FA-treated compared with the untreated group (Fig. 6F). Thus, exogenous FA treatment increased Tex^STEM^ numbers and percentages, while also decreasing Tex^INT^, increasing PD1 expression and compromising viral control. The increase in Tex^STEM^ upon FA treatment may be attributed to their enhanced capacity to uptake and metabolize medium and long-chain FA.

**Fig 6.**
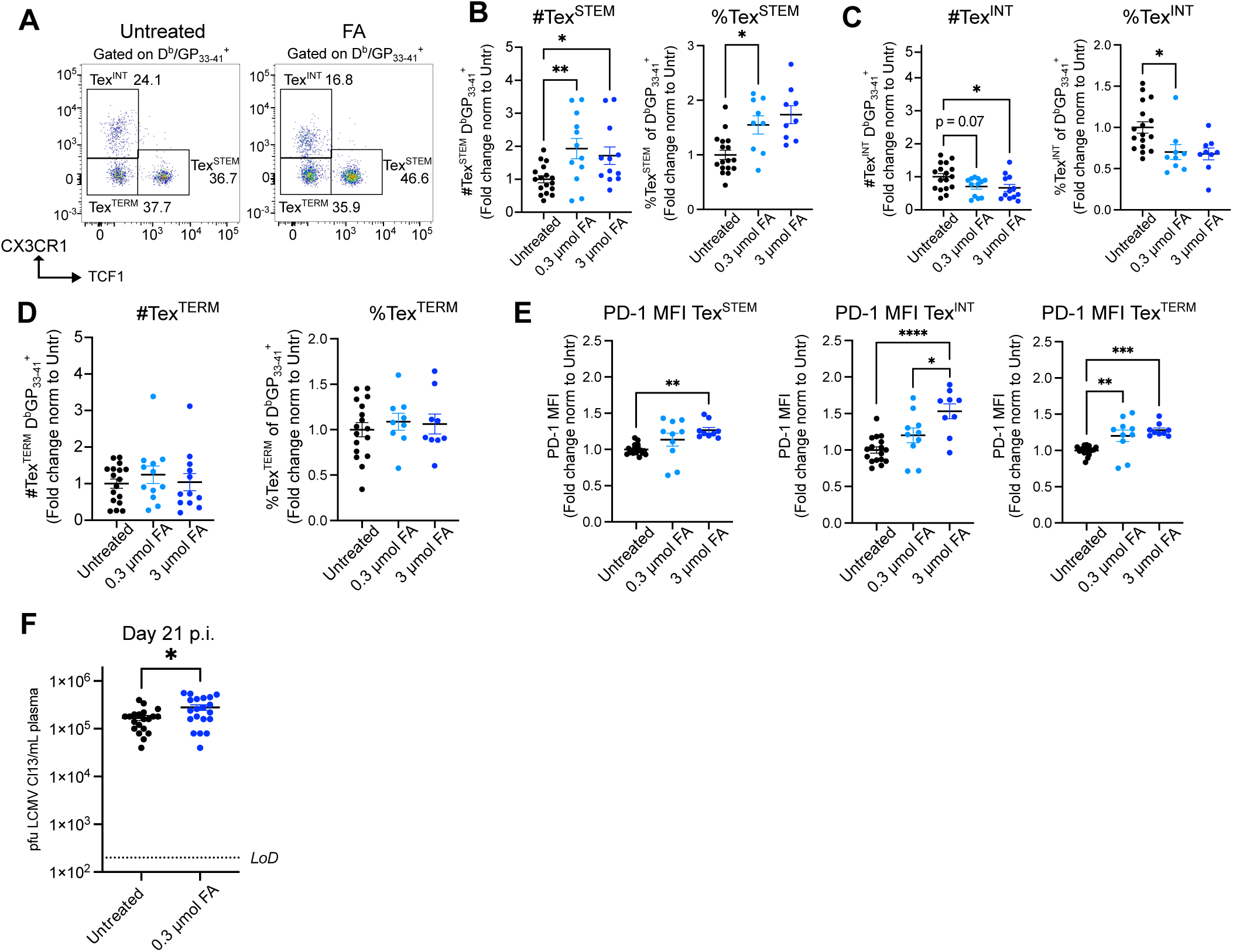
*In vivo* FA administration increased Tex^STEM^ while reducing Tex^INT^ and impairing viral control during LCMV Cl13 infection. C57BL/6 mice were infected with LCMV Cl13 and treated i.p. with either 0.3 mmol (A-F) or 3 mmol (A-E) of FA (1:1 lauric and palmitic acid mix) or PBS every 12 hours from days 16 to 20 p.i.. (A-F) Splenocytes (A-E) or blood (F) were obtained at day 21 p.i.. (A) Flow cytometry showing CX3CR1 and TCF1 expression in CD8^+^ D^b^/GP_33-41_ tetramer^+^ cells. (B-D) Fold change of numbers (left) or percentages (right) of CD8^+^ D^b^/GP_33-41_ tetramer^+^ Tex^STEM^ (LY108^+^ CX3CR1^-^) (B), Tex^INT^ (LY108^-^ CX3CR1^+^) (C) or Tex^TERM^ (LY108^-^ CX3CR1^-^) (D) normalized to the average from untreated mice at day 21 p.i.. (E) Mean fluorescence intensity (MFI) of PD-1 in CD8^+^ D^b^/GP_33-41_ tetramer^+^ Tex^STEM^ (left), Tex^INT^ (middle) or Tex^TERM^ (right) normalized to the average from untreated mice at day 21 p.i.. (F) Plasma viral titers from FA treated mice at day 21 p.i.. (B-F) Averages ± SEM. (A-E) Data are combined across three independent repeats with n=4-5 mice/group. (F) Data are combined across four independent repeats with n=4-8 mice/group. (B-E) One-way ANOVA with Tukey’s correction. (F) Individual two-tailed Student’s t-test. *p<0.05, **<0.01, ***<0.001, ****<0.0001.

## Discussion

While immune responses are initiated by the direct recognition of pathogens, they are also shaped by factors in the surrounding environment, including systemic metabolites that reach lymphoid and non-lymphoid tissues through circulation(61). The dynamic changes in the systemic metabolome during acute vs. chronic viral infections and the specific ways these metabolites influence immune responses are, however, only beginning to be understood(62, 63). In the present study we used untargeted metabolomics to compare the systemic chemical environments induced early and late following acute vs. chronic LCMV infection.

It was noteworthy that most chemical adaptations observed during LCMV Cl13 infection were transient and did not associate with sustained viral loads; instead, they resembled metabolic changes typically observed during fasting(47). Combined with the profound short-term anorexia that we and others have reported in the first week following LCMV Cl13 infection(45, 46), these findings suggest that the early metabolic shifts during a persistent infection are partially driven by a temporary reduction in food intake, explaining their transient nature. Our comparison with the metabolome in fasted humans(47) also highlighted the conservation of the metabolomic changes that emerge upon reduced food intake across species, whether due to infection or starvation.

While many of the metabolite shifts that we detected likely convey important signals to immune cells, one notable metabolic signature early after LCMV infection was the significant elevation of medium- and long-chain FA and acylcarnitines. This elevation is consistent with previously reported plasma elevation of free FA and white adipose tissue lipolysis after LCMV Cl13 infection(45). We validated and extended these findings by demonstrating that plasma FA elevation and white adipose tissue lipolysis occur in both LCMV ARM and Cl13 infections, although the degree was significantly more pronounced in the latter. This is consistent with previous reports documenting adipose tissue lipolysis in acute bacterial infection models(64) and cancers(65, 66). In such cases, however, cytokines such as TNF-α, IL6 and/or IFN-γ have been identified as causative factors(67–69). In contrast, in the LCMV Cl13 infection model, where weight loss appears associated with lipolysis, these cytokines were ruled out as the mediators responsible for the observed weight reduction(45). Instead, CD8 T cells were implicated in adipose tissue lipolysis, as evidenced by the attenuated weight loss and reduced free FA previously reported in T cell-depleted or deficient LCMV-infected mice (45). These findings are further supported by our observations of decreased long-chain FA and acylcarnitines in addition to reduced adipocyte HSL phosphorylation upon CD8 T cell depletion. Notably, elevation of free long-chain FA and acylcarnitines is also a signature of fasting(47), which is known to induce lipolysis(70). This supports a model in which CD8 T cell-induced anorexia contributes to WAT lipolysis and the rise in systemic long-chain FA related species early after LCMV Cl13 infection. Notably, this model does not exclude the possibility that other mechanisms may also contribute to lipolysis and FA elevation in acute and/or chronic infections.

The pronounced elevation of medium- and long-chain FA after LCMV infection prompted us to explore their impact on CD8 T cell responses, illustrating the value of our metabolomics dataset. We first revealed that virus-specific CD8 T cells from LCMV Cl13-infected mice exhibited higher neutral lipid accumulation as well as medium- and long-chain FA uptake than their counterparts from LCMV ARM-infected mice, which may be related to an adaptation to the greater availability of systemic FA and lower glucose that we observed in the chronic setting. In addition, we found that within the exhausted CD8 T cell population in LCMV Cl13 infection, Tex^STEM^ cells were particularly adept at accumulating, up-taking and metabolizing FA to fuel oxidative phosphorylation, aligning with previous observations of enhanced oxidative phosphorylation in this Tex subset(31). Consistently, reintroduction of FA late after LCMV Cl13 infection, when endogenous FA are no longer elevated, increased Tex^STEM^ mitochondrial fitness. These findings raised the possibility that the elevation in medium- and long-chain FA that naturally takes place early after LCMV Cl13 infection may fuel the metabolism of Tex^STEM^ present during this period. This possibility would be consistent with the many similarities between Tex^STEM^ and memory CD8 T cells(30) and the observation that FAO fuels memory CD8 T responses(71, 72). It is important to note, however, that unlike what we observed in Tex^STEM^, circulating memory CD8 T cells acquire minimal amounts of extracellular FA(73), suggesting that these two long-lived CD8 T cell populations potentially use different FA sources to fuel FAO. Conversely, tissue resident memory T cells use various fatty acid binding proteins to uptake lipids and optimize their long-term maintenance across different tissues(74, 75).

Interestingly, we found that short-term FA administration *in vivo* also increased the numbers and percentages of Tex^STEM^, but not other Tex subsets. This numerical increase was particularly striking given that FA exposure appeared to up-regulate expression of genes related to apoptosis while down-regulating cell-cycle associated genes in Tex^STEM^. While the mechanisms underlying this effect remain to be elucidated, a recent report showed that enhancing FAO via inhibition of acetyl-CoA carboxylase activity in tumor-infiltrating lymphocytes (TILs) upregulated stemness markers, including TCF1 expression(76). Thus, the increased Tex^STEM^ numbers following FA exposure are likely a consequence of heightened FAO, which in turn could have promoted a stemness state while disfavoring Tex^STEM^ proliferation and differentiation. Since Tex^STEM^ are crucial for generating effector-like Tex^INT^ cells(30), which ultimately control viral replication(28, 29), it is tempting to speculate that this putative attenuation of Tex^STEM^ differentiation could have contributed to the reduced numbers of Tex^INT^ cells and higher viral titers observed after FA administration during late LCMV Cl13 infection. It is, however, also possible that FA treatment could have directly or indirectly affected Tex^INT^, independently of an effect on Tex^STEM^ differentiation, and/or that the increased viral burden resulted from an independent effect of FA on the observed PD-1 up-regulation. Further studies are needed to harness the benefits that FA exposure can confer on Tex^STEM^ metabolism and numbers without compromising Tex^INT^ and viral control.

From an evolutionary perspective, and considering that evolution has been shaped by numerous periods of starvation(77), a system where stem-like cells, such as Tex^STEM^, become the most avid for medium- and long-chain FAs, the predominant nutrient in the absence of food consumption, likely offers significant advantages. This mechanism would enable stem-like cells to outcompete their differentiated progeny for the limited available nutrients, ensuring their survival and metabolic fitness so they can be preserved for replenishment of progeny when normal food intake resumes. In addition, stem-like cells may dwell longer within bone marrow to enhance their survival during nutrient shortage, as shown for memory CD8 T cells(78). On the other hand, the fact that CD8 T cell responses drive the natural elevation of FA, which we found are capable of limiting the numbers of effector-like Tex^INT^, suggests a potential negative feedback loop, likely shaped by evolutionary pressure to limit CD8 T cell-induced immunopathology upon infection with a rapidly replicating virus.

Finally, it is noteworthy that the tumor microenvironment is also rich in lipid species(79–84). However, unlike chronic LCMV infection, FA elevation in tumors is long-term(79–84). Prolonged FA elevation in the tumor microenvironment may predispose it to lipid oxidation, with oxidized lipids taken up by TILs and leading to peroxidation and dysfunction(79). Finally, TILs also exhibit a reduced capacity to metabolize long-chain FAs, which further contributes to their functional exhaustion, although this study did not distinguish the effect on different Tex subsets(83). Further studies are necessary to investigate whether Tex^STEM^ from tumor settings are also preferentially capable of up-taking, metabolizing, and benefiting from long-chain FA present in the tumor microenvironment.

Overall, our work uncovers a distinct systemic metabolic landscape in acute vs. chronic viral infections, marked by significant shift in nutrient availability that is mostly transient and more pronounced in the latter condition. The short-term elevation of medium- and long-chain FA, along with their preferential uptake and metabolism in Tex^STEM^, as well as the exclusive increase in Tex^STEM^ upon FA administration, exemplifies the influence that changes in systemic metabolites could have on immune responses. Our comprehensive metabolome profiling offers a valuable resource for the scientific community and encourages further exploration on the role of individual metabolites in shaping antiviral immunity during acute and chronic infections.

## Methods

### Mice, infections and treatments

Six to ten-week-old female C57BL/6 mice were purchased from The Jackson Laboratory and housed under specific-pathogen-free conditions at the University of California, San Diego. Mice were infected intravenously (i.v.) with 2×10^6^ plaque forming units (PFU) of LCMV ARM or Cl13. Viral stocks were grown, identified and quantified as reported previously(85). Viral loads were determined by standard plaque assay in VERO cells of plasma samples. For CD8 T cell depletion studies, mice were intraperitoneally injected with anti-CD8α (53-6.72, BioXcell) or IgG2a (2A3, BioXcell) antibodies on day -2, -1, and 5 p.i. (250 µg/mouse) as well as on the day of infection (200 µg/mouse). For FA administration studies, mice were intraperitoneally injected with 0.3 or 3.0 μmol of a 1:1 mixture of lauric and palmitic acid dissolved in castor oil twice daily starting at the onset of the dark cycle at day 16 p.i. and concluding at the onset of the light cycle at day 20 p.i. For acute starvation experiments, mice were food-restricted for a total of 48 hours. Mice were maintained in a closed breeding facility where they were housed in ventilated cages containing HEPA filters. Mouse handling conformed to the requirements of the National Institutes of Health and the Institutional Animal Care and Use Guidelines of UCSD.

### Sample collection for metabolomic studies

For metabolomic studies, mice were bled via submandibular bleed and blood was collected with heparinized capillaries (ThermoFisher). To obtain plasma, blood was then spun down at 10,000 rpm for 10 min at 4°C and plasma aliquoted and placed at -80°C.

### Lymphocyte isolation from spleen

Spleens were harvested and digested with 1mg/mL collagenase D for 20 minutes at 37°C and mechanically dissociated through a 100µm filter to make a single cell suspension. Cells were then centrifuged at 1500 rpm at 4°C for 5 minutes. Red blood cells were lysed by incubating pellets in 1mL ammonium-chloride-potassium (ACK) lysis buffer (150mM NH_4_Cl, 10mM KHCO_3_, 0.1mM Na_2_EDTA in deionized water, at pH 7.4) for 5 minutes at room temperature (RT). The lysis was quenched by the addition of 10mL of 1× PBS + 3% fetal bovine serum (FBS). Cells were pelleted as above. Pellets were resuspended in 2mL complete Roswell Park Memorial Institute (RPMI, Gibco), which consisted of RPMI supplemented with 10% heat-inactivated Fetal Bovine Serum (FBS, Atlanta Biologicals), 1% Penicillin/Streptavidin (P/S, BioWhittaker), 1 mM Sodium Pyruvate (Na-pyr, ThermoFisher Scientific), 1mM L-Glutamine (L-Gln, BioWhittaker), and 20 mM HEPES 2 (ThermoFisher Scientific). Absolute lymphocyte numbers were determined by forward scatter/side scatter gating with a Guava Easycyte automated cell counter (MilliporeSigma, MA).

### Western blot in adipose tissue

Inguinal WAT was bead-homogenized in the presence of RIPA buffer (ThermoFisher Scientific) supplemented with phosphatase and protease inhibitors (Roche Applied Science). All lysates were cleared by centrifugation at 12,000 rpm for 30 min at 4 °C and total protein quantified by standard Bradford assay. Protein extracts were resolved by 10% SDS-PAGE, transferred to PVDF membranes (EMD Millipore) and blocked for 1hr at room temperature (RT) with 3% non-fat dry milk in PBS containing 0.1% Tween-20. Membranes were probed with the primary antibodies anti-GAPDH (14C10) Rabbit mAb (1:5000), anti-HSL Rabbit mAb (1:5000) or anti-phospho-HSL (Ser660) Rabbit mAb (dilution 1:3000), overnight at 4°C, followed by incubation with HRP-conjugated antibody (1:5000) for 1 hour at RT. Blots were visualized using SuperSignal West Pico PLUS Chemiluminescent Substrate (Thermo Scientific) with an Odyssey CLx Imager (LI-COR Biosciences). All antibodies obtained from Cell Signaling Technologies.

### Quantification of adipocyte area

Visceral WAT (adjacent to the reproductive tract) was fixed in 10% formalin for 24 hours, followed by 2 washes with distilled water and dried in 70% ethanol. Tissues were embedded in paraffin, sectioned and stained with hematoxylin and eosin by the Tissue Technology Shared Resources (TTSR) at the Moore’s Cancer Center, La Jolla, CA. Adipocyte area quantification was done with ImageJ(86) on images at a magnification of 10X. A macro was created that converted images to 8-bit, adjusted the threshold, switched to a dark background, applied watershed and analyzed particles. We quantified 3 different fields per mouse that consistently contained ∼50-150 adipocytes each. Data shown correspond to pooled raw adipocyte areas from all fields and all mice within each experimental group.

### Quantification of glucose and plasma free FA

Absolute blood concentrations of blood glucose were quantified with a standard glucometer (Germaine Laboratories). Plasma samples were analyzed with the Free Fatty Acid Assay Kit (Abcam) per manufacturer’s instructions, following the fluorometric detection method.

### Quantitative PCR

For quantification of RNA transcripts in the WAT, tissues were collected, snap frozen, and stored at -80°C. Thawed tissues were bead homogenized in RLT buffer (Qiagen) and centrifuged for 10 min at 10,000 rpm. Supernatants were collected for total RNA extraction using RNeasy kits (Qiagen), digested with DNase I using the RNase-free DNase set (Qiagen) and reversed transcribed into cDNA using M-MLV Reverse Transcriptase (Promega). The expression of genes was quantified, in triplicates, using Fast SYBR Green Master Mix (ThermoFisher Scientific) and the CFX96 Touch Real-Time PCR Detection System (Bio-Rad). Relative transcript levels were normalized against murine GAPDH. Graphs depicting qRT-PCR analysis represent biological replicates of individual mice from one representative experiment.

### Flow Cytometry

Cells were stained at a maximal concentration of 2×10^7^ cells/mL in FACS buffer (1× PBS + 3% fetal bovine serum (FBS). For surface staining, cells were initially stained with a fixable viability dye (Tonbo Ghost dye) in 1×PBS for 10 min at 4°C followed by staining with the MHC-I tetramer H2-D^b^/GP_33-41_-BV421 provided by NIH Tetramer Core Facility (Atlanta, GA) in FACS buffer for 1hr at RT, followed by staining for 20 min at 4°C with remaining surface antibodies. For intranuclear staining, cells were fixed using the Foxp3 Transcription Factor Staining Buffer Set Kit (ThermoFisher Scientific), following the vendor’s recommendations. Alternatively, for intracellular staining after *ex vivo* peptide stimulations and neutral lipid stains, cells were fixed in 1% PFA for 20 minutes at 4°C. After fixation, cells were stained with antibodies in 10x Permeabilization Buffer (ThermoFisher Scientific), diluted 1:10 in water, for 1 hour at RT. Antibodies used in this study were purchased from ThermoFisher Scientific (Waltham, MA), BD Biosciences, or Biolegend (San Diego, CA).

The following antibodies were used to stain single cell suspensions prepared from murine spleens or *ex vivo* cultures: CD44 (IM7), CD8α (53-6.7), CX3CR1 (SA011F11), IFNγ (XMG1.2), LY108 (330-AJ), PD-1 (29F.1A12), TCF1 (C63D9) and TIM3 (RMT3-23). Cells were acquired using a ZE5 Cell Analyzer (Bio-Rad). Flow cytometric data were analyzed with FlowJo software v10.

### *Ex vivo* T cell stimulation

Splenocytes were cultured at 1×10^7^ cells/ml in round-bottom 96-well plates for 5 hr in complete RPMI supplemented with Brefeldin A (1 μg/mL; Sigma) and MHC class-I-restricted LCMV GP_33– 41_ peptide (1 μg/mL, all >99% pure; Synpep). Cells were then stained with a fixable viability dye (Tonbo Ghost Dye) and cell surface antibodies as described above. After fixation in 1% PFA, cells were stained with an anti-IFNγ antibody in Permeabilization Buffer (ThermoFisher Scientific) for 30 min at RT. Unstimulated controls in which cells were cultured without peptide were performed in parallel.

### Fatty acid uptake, neutral lipid content and mitochondrial staining assays

Fatty acid uptake and neutral lipid content were evaluated as described in (79). Briefly, for lipid uptake assays, isolated cells were incubated with 0.5μg/mL C1-BODIPY 500/510 C12 (ThermoFisher) or BODIPY FL C16 (ThermoFisher) in 1×PBS for 20 min at 37°C. After incubation, cells were washed in FACS buffer and stained with cell surface antibodies as described above, without any fixation or permeabilization. Cells were acquired using a ZE5 Cell Analyzer (Bio-Rad) immediately after staining. For neutral lipid content, isolated cells were stained with cell surface antibodies and fixed with 1% PFA for 20 min at 4°C. Cells were then permeabilized and stained using BODIPY 493/503 (ThermoFisher) at a final concentration of 250 ng/mL in Permeabilization Buffer (ThermoFisher Scientific). Cells were acquired using a ZE5 Cell Analyzer (Bio-Rad). Mitochondrial polarization was evaluated as described in (58), (79). Briefly, cells were incubated with 10nM MitoTracker Red CMXRos (ThermoFisher) and 10nM MitoTracker Green FM (ThermoFisher), concurrently, for 15 min. After staining, cells were washed in 1X PBS, followed by surface marker staining as described above. Cells were not fixed or permeabilized. The ratio of mitochondrial membrane potential to mitochondrial mass was calculated by dividing the mean fluorescent intensity (MFI) of MitoTracker Red CMXRos (membrane potential) by the MFI of MitoTracker Green FM (mass). Cells were acquired using a ZE5 Cell Analyzer (Bio-Rad) immediately after staining.

### Cell sorting

Isolated splenocytes were enriched using EasySep^TM^ Mouse Streptavidin RapidSpheres^TM^ Kit (Stemcell Technologies) per the manufacturer’s instructions. Biotin conjugated anti-CD19 (1D3), anti-B220 (RA3-6B2) and anti-CD4 (RM4-5) were used. Cells were stained with propidium iodide (PI) (Sigma Aldrich) and Tex subsets FACS-purified by the following definitions: Tex^STEM^ (PI^-^, CD8^+^, PD1^+^, CD44^+^, CX3CR1^-^ LY108^+^); Tex^INT^ (PI^-^, CD8^+^, PD1^+^, CD44^+^, CX3CR1^+^, LY108^-^); Tex^TERM^ (PI^-^, CD8^+^, PD1^+^, CD44^+^, CX3CR1^-^, LY108^-^). Cell sorting was performed on a BD FACSAria II (BD Biosciences) or BD FACSAria Fusion (BD Biosciences) flow cytometer.

### *Ex vivo* T cell culture

FACS-purified Tex subsets were cultured at 5×10^5^ cells/mL in round-bottom 96-well plates for 8 hours in complete RPMI media supplemented with 55 μM beta-mercaptoethanol, a cocktail of the MHC class-I-restricted peptides (GP_33-41_, GP_276-286_, and NP_396-404_), and 200μM of either BSA or BSA-palmitate, with a final volume of 200 μL per well. After culture, cells were harvested for RNA sequencing (detailed below).

### Metabolic Flux Assays

To analyze oxygen consumption rate (OCR), Seahorse XFp plates and cassettes (Agilent) were prepared per the manufacturer’s instructions. Tex subsets were isolated by FACS and resuspended in RPMI Seahorse Assay Medium with the addition of 2.5 mM glucose, 0.5 mM carnitine, and 5mM HEPES. 2×10^5^ cells/well were plated into a Seahorse XF HS Miniplate (Agilent). Cells were then rested for 1 hour at 37°C without CO_2_. Immediately before initiating the assay, prewarmed BSA or palmitate conjugated to BSA was added to the corresponding wells, for a final concentration of 200μM. Assays were performed on a Seahorse XF HS Mini analyzer. A mitochondrial stress test assay on the XF-HS Mini was optimized within the ranges provided by the manufacturer and performed using the Mitochondrial Stress Test kit (Agilent). In-well concentrations for each drug at the time of injection were Oligomycin (1.5 μM), FCCP (2 μM) and Antimycin/Rotenone (0.5 μM). Basal OCR was calculated as the difference between OCR prior to Oligomycin addition and OCR after Antimycin/Rotenone. Spare respiratory capacity was calculated as the difference between OCR prior to the addition of Oligomycin and OCR measured after FCCP addition. ATP production related oxygen consumption was calculated as the difference between OCR prior to the addition of Oligomycin and the OCR after Oligomycin addition. Mitochondrial independent oxygen consumption was calculated as the OCR value after the addition of Antimycin/Rotenone.

### Metabolite extraction and UPLC-Q-TOF mass spectrometry analysis

Plasma samples were extracted by adding 100% methanol spiked with 2 µM sulfamethazine to a final concentration of 80% methanol. Samples were vortexed for two minutes and then placed at - 20°C for 20 min to aid protein precipitation. Samples were then centrifuged at 15,000 and 80% of the supernatant was placed into a 96 well plate. The plates were then lyophilized using a CentriVap Benchtop Vacuum Concentrator (Labconco) and stored at -80°C. At the time of analysis, the lyophilized samples were resuspended in 50% methanol spiked with 1µM sulfadimethoxine. The metabolomic extracts were analyzed using an ultra-high-performance liquid chromatography (UHPLC) system (Vanquish, Thermo Scientific) coupled to a quadrupole-Orbitrap mass spectrometer (Q Exactive, Thermo Scientific). Chromatographic separation was performed using a Kinetex C18 1.7 μm, 2.1 mm x 50 mm column (Phenomenex) at a flow rate of 0.5 mL/min and held at a temperature of 40°C. The mobile phase consisted of solvent A, 100% LC-MS grade water with 0.1% formic acid (v/v) and solvent B, 100% LC-MS grade acetonitrile with 0.1% formic acid (v/v). The chromatographic elution gradient was: 0.0–1.0 min, 5% B; 1.0–9.0 min, 5–100% B; 9.0-11.0 min, 100% B; 11.0-11.5 min, 100-5% B; 11.5-12.5 min, 5% B. Heated electrospray ionization parameters were: spray voltage, 3.5 kV; capillary temperature, 268.0°C; sheath gas flow rate, 52.0 (arb. units); auxiliary gas flow rate, 14.0 (arb. units); auxiliary gas heater temperature, 435.0°C; and S-lens RF, 50 (arb. units). MS data was acquired in positive mode using a data dependent method with a resolution of 35,000 in MS1 and a resolution of 17,000 in MS2. An MS1 scan from 100-1500 m/z was followed by an MS2 scan, using collision induced dissociation, of the five most abundant ions from the prior MS1 scan.

### LC-MS data processing and availability

Q Exactive .raw files were converted to .mzXML format using ProteoWizard’s msConvert (87). MS1 feature finding for all samples and controls was done using the open-source software MZmine version 2.40(88) with the following parameters: 1) Mass detection > Centroid > Noise level 3.0E5; 2) Chromatogram builder: minimum time span 0.01 min, minimum height 3.0E5, m/z tolerance 10 ppm; 3) Deconvolution: m/z range for MS2 scanning 0.01, RT range MS2 0.25. For deconvolution we used Local Minimum Search with the following parameters: chromatographic threshold 0.01%, search minimum RT 0.2 min, minimum relative height 0.01%, minimum absolute height 3.0E5, minimum ratio of peak/edge: 3, peak duration range 0.05-0.5 min. 4) Isotope grouper: m/z tolerance 10 ppm, RT 0.25, maximum charge: 4. 5) Join aligner: m/z tolerance 10 ppm, weight for m/z: 90, RT tolerance 0.3 min, weight for RT 10. The resulting matrix was exported as a .csv file. Features appearing in blanks, heparin controls, and washes were eliminated from all samples, followed by normalization to the resuspension solvent internal standard, sulfadimethoxine, and followed by an additional normalization to each sample’s total area, for a final representation of relative abundance of all molecular features. The resulting matrix was converted to biom format and uploaded to Qiita(89) along with corresponding metadata formatted as proposed by the Earth Microbiome portal(90) under study ID: 11043 for downstream analysis. All MS (.raw, .mzXML, etc) were uploaded to UCSD MassIVE (https://massive.ucsd.edu/)(91) and are publicly accessible under IDs: MSV000081343 (contains plasma samples from uninfected and LCMV-infected mice at day 8 and 20 post-infection) and MSV000084904 (contains plasma samples from isotype and anti-CD8-treated, LCMV-infected mice, and uninfected mice fed ad libitum or fasted).

### Metabolite Identification

Molecular networking was performed using the ‘Data Analysis’ workflow on Global Natural Products Social Molecular Networking portal (GNPS; https://gnps.ucsd.edu/)(42). Note that annotations obtained in this way are MSI level 2 or 3 according to the 2007 metabolomics standards initiative(43). A molecular network was created using the online workflow at GNPS. The data was filtered by removing all MS/MS peaks within +/- 17 Da of the precursor m/z. MS/MS spectra were window filtered by choosing only the top 6 peaks in the +/- 50Da window throughout the spectrum. The data was then clustered with MS-Cluster with a parent mass tolerance of 0.02 Da and a MS/MS fragment ion tolerance of 0.02 Da to create consensus spectra. Further, consensus spectra that contained less than 2 spectra were discarded. A network was then created where edges were filtered to have a cosine score above 0.7 and more than 4 matched peaks. Further edges between two nodes were kept in the network if and only if each of the nodes appeared in each other’s respective top 10 most similar nodes. The spectra in the network were then searched against GNPS’ spectral libraries. The library spectra were filtered in the same manner as the input data. All matches kept between network spectra and library spectra were required to have a score above 0.7 and at least 4 matched peaks. Link to analyses shown on Fig. 1 and 2 are provided. Day 8 – uninfected, ARM, Cl13: https://gnps.ucsd.edu/ProteoSAFe/status.jsp?task=93dc1e3f1dee4218939ebdc04264461c Day 20 – uninfected, ARM, Cl13: https://gnps.ucsd.edu/ProteoSAFe/status.jsp?task=6231f4e1ae024169b60d7d3633503f03 Ad libitum vs fasting, Isotype vs anti-CD8: https://gnps.ucsd.edu/ProteoSAFe/status.jsp?task=5c5b89870f9d488cb96ffca88ef32282 Spectral libraries staticlibs/NIST_17 were used.

### Metabolomics data analysis

Plasma metabolomics data were analyzed on Qiita under analysis IDs 31008 and 31009. We calculated beta-diversity by the Jaccard similarity index (‘beta’ command) followed by PCoA analysis (‘pcoa’ command) and significance assessment by PERMANOVA with 999 permutations (‘beta_group_significance’ command).

Pairwise comparisons between Un, ARM-infected and Cl13-infected mice were all performed on Metaboanalyst, under the ‘Statistical Analysis’ module(92). Significance was determined by Wilcoxon rank test with false discovery rate (FDR) below 0.05 unless otherwise indicated. Heatmaps were generated with Metaboanalyst using ANOVA using FDR below 0.05 showing top 200 differential metabolites based on f-value. Volcano plots were generated with Metaboanalyst using Student t-test and FDR below 0.05. Venn diagrams were constructed via (http://bioinfogp.cnb.csic.es/tools/venny/). Metabolic pathway prediction was done by using mummichog v.1.0.9 with default parameters(41). The original matrix compatible with statistical analyses and containing detected m/z, retention-times and normalized intensities that was used in(47) was used for metabolic pathway prediction in Fig. 2J.

### RNA Sequencing

For RNA sequencing (RNAseq) studies, cells were obtained after *ex vivo* cultures or *in vivo* FA treatment as described above. Samples from independent experiments were kept at -80C in RLT buffer from RNeasy mini kits (Qiagen) containing 10μL β-mercaptoethanol per mL RLT (per manufacturer’s instructions) until RNA extraction. Total RNA was prepared from 50,000-75,000 cells. RNA was processed by the UCSD IGM genomics center to generate an RNAseq library using a TruSeq RNA Library Prep Kit v2 (Illumina). Raw sequencing data were aligned to mouse genome mm10 (Genome Reference Consortium GRCm38) by using Spliced Transcripts Alignment to a Reference (STAR) aligner106 with the following parameters: “--outFilterType BySJout -- outFilterMultimapNmax 20 --alignSJoverhangMin 8 --alignSJDBoverhangMin 1 -- outFilterMismatchNmax 999 --outFilterMismatchNoverReadLmax 0.04 --alignIntronMin 20 -- alignIntronMax 1000000 --alignMatesGapMax 1000000”. Downstream analyses and PCA plots were generated with R 4.3.3(93) and Bioconductor 3.19(94). Differentially expressed genes (DEGs) were determined by DEseq2(95) using a log2 fold change of ≥0.5 with a p-adjusted value of ≤0.05 as the threshold. Pathway analyses were done with ShinyGo(96) and PathFindR(97).

### Statistical analysis

Statistical analysis methods for metabolomics and RNAseq data are detailed in the respective methods section. For non-omics data, unpaired two-tailed Student’s t-test was used to compare two independent groups, or paired t-test when samples were derived from the same animal or same pool of cells from different animals. If variances were not equal by F-test, data were tested using the non-parametric Mann Whitney-U test. Significant differences among three groups were determined based on one-way ANOVA with Tukey’s multiple comparison correction or, in the case of unequal variances, non-parametric Kruskal-Wallis test with Dunn’s multiple comparison correction. Analyses of non-omics data were performed using GraphPad Prism v10.

## Data Availability

Metabolomics datasets reported in this study are available as described above.

RNAseq datasets reported in this study will be uploaded to Gene Expression Omnibus (GEO) prior to publication.

## Code Availability

All software used to perform these analyses are publicly available. Software tools are listed in the results and/or methods sections.

## Acknowledgements

The authors are grateful to members of the Zúñiga lab, who provided valuable discussions and critical evaluation of this manuscript. We thank personnel at the UCSD Animal Care Program, IGM Genomics Center sequencing core, La Jolla Institute flow cytometry core, Sanford Burnham Prebys flow cytometry core, and Moore’s Cancer Center Tissue Technology Shared Resources (TTSR). We are grateful to Cody Fine, Mateo Espinoza and Mitra Banihassan (UCSD) for technical assistance with flow cytometry experiments. This work was additionally made possible by the UC San Diego Stem Cell Program and a CIRM Major Facilities grant (FA1-00607) to the Sanford Consortium for Regenerative Medicine. This publication includes data generated at the UCSD Human Embryonic Stem Cell Core Facility, using the FACS Aria Fusion and FACS Aria II Flow Cytometry Sorters, and at the UC San Diego IGM Genomics Center utilizing an Illumina NovaSeq 6000 that was purchased with funding from a National Institutes of Health SIG grant(#S10OD026929).

This study was supported by NIH grants AI081923, AI135314, and AI113923 as well as a UCSD Center for Microbiome Innovation seed grant to E. I. Zúñiga. K. R. Kazane was partially supported by the UCSD T32 Training Grant.

## Author Contributions

K. R. Kazane designed, performed, analyzed and interpreted T cell experiments, helped with metabolome data analysis and wrote the manuscript. L. Labarta-Bajo designed, performed, analyzed and interpreted metabolomics experiments and adipose tissue lipolysis and fasting related experiments, and aided in writing of corresponding parts in the manuscript. F. Vargas and K. C. Weldon performed and analyzed metabolomics experiments. K. Liimatta aided with T cell experiments. D. R. Zangwill analyzed RNA sequencing experiments and helped with metabolome data analysis. P. C. Dorrestein supervised metabolomics analysis. E. I. Zúñiga conceived and supervised the project, designed and interpreted experiments, and wrote the manuscript. All authors revised and approved the manuscript.

## Figures

**Fig S1.**
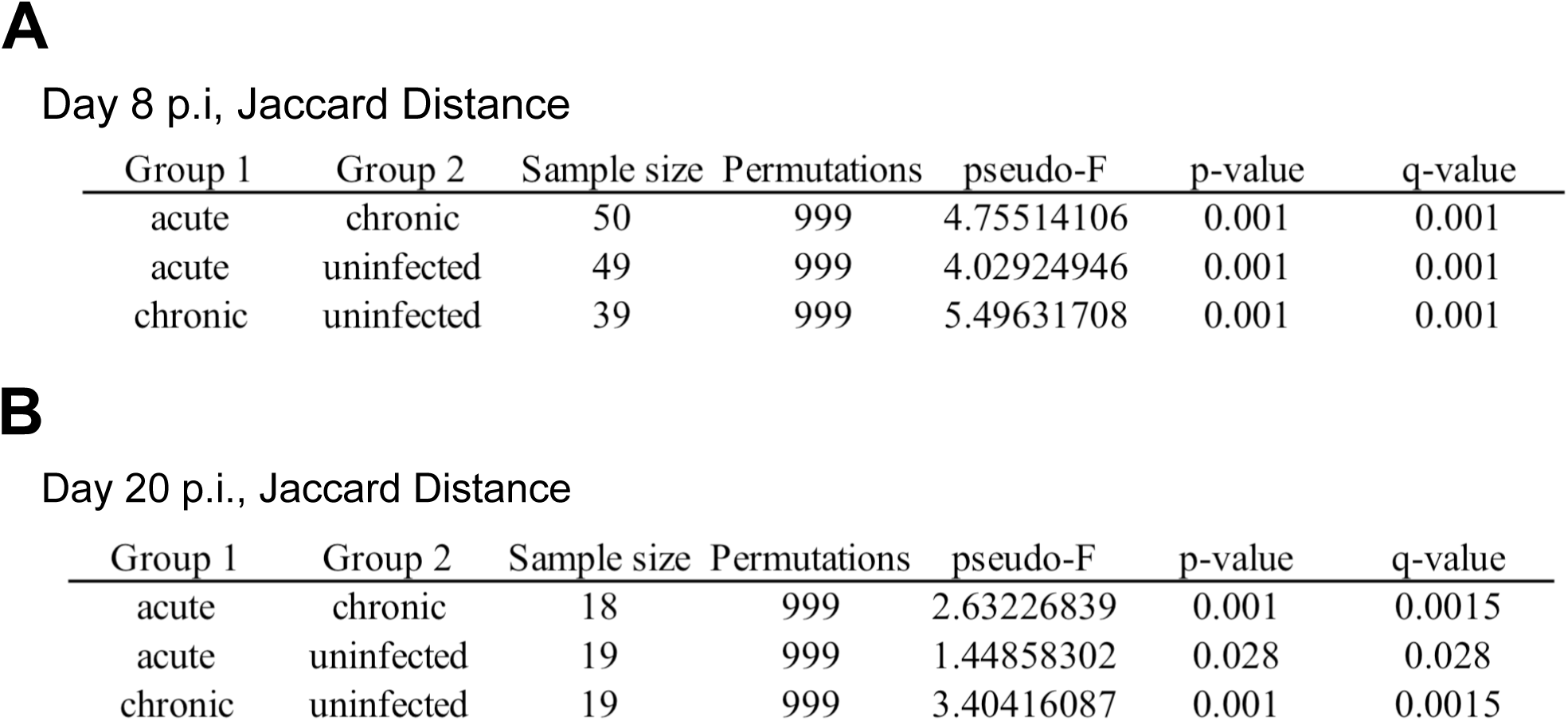
Untargeted metabolomics analysis revealed that uninfected, ARM- and Cl13-infected mice form significantly distant clusters at days 8 and 20 p.i. C57BL/6 mice were infected with LCMV ARM, Cl13 or left uninfected (Un) and untargeted metabolomics was performed on plasma from day 8 and 20 p.i.. PCoA plots with Jaccard distance metric were generated and beta group significance was calculated by Permanova (999 permutations) at day 8 p.i. (A) and 20 p.i. (B). (A) Data show 2 pooled experiments with n=10-20 mice/group on day 8 p.i. and (B) 1 experiment with n=9-10 mice/group on day 20 p.i..

**Fig S2.**
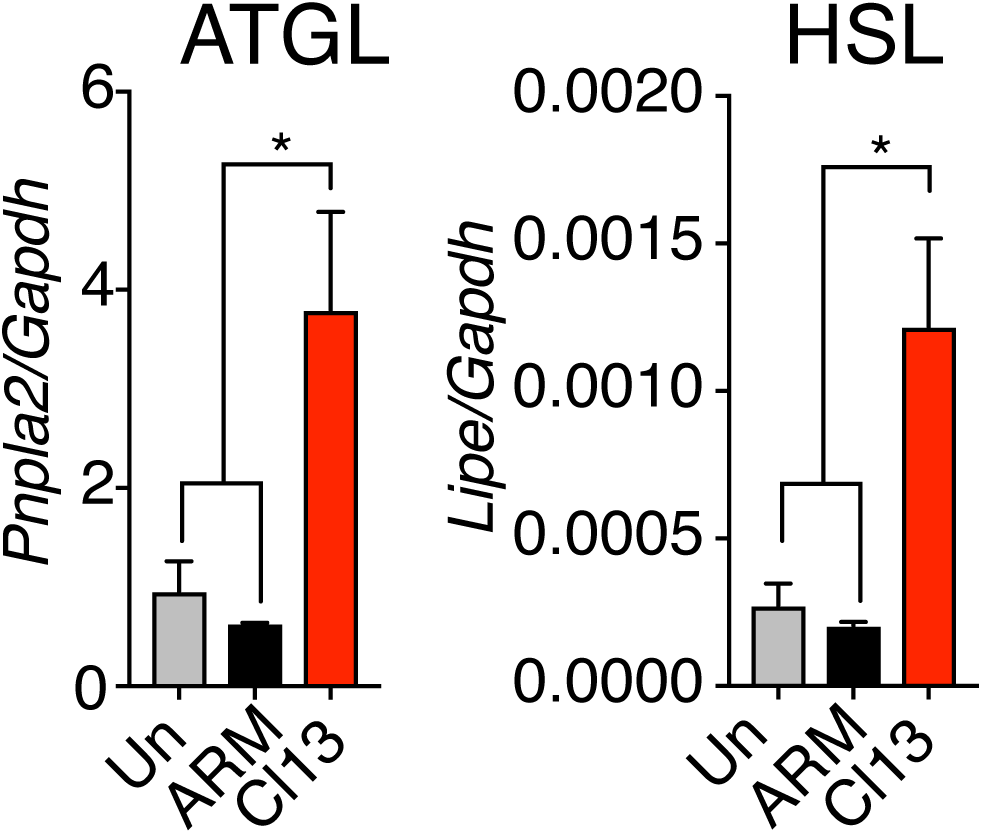
LCMV Cl13 infection increased lipase transcripts in white adipose tissue. C57BL/6 mice were infected with LCMV ARM, Cl13 or left uninfected (Un). *Pnpla2* and *Lipe* mRNA levels relative to *Gapdh* in visceral white adipose tissue at day 9 p.i.. Representative of 2 independent experiments with n=3-5 mice/group. Averages ± SEM. One-way ANOVA with Tukey’s correction. *p<0.05.

**Fig S3.**
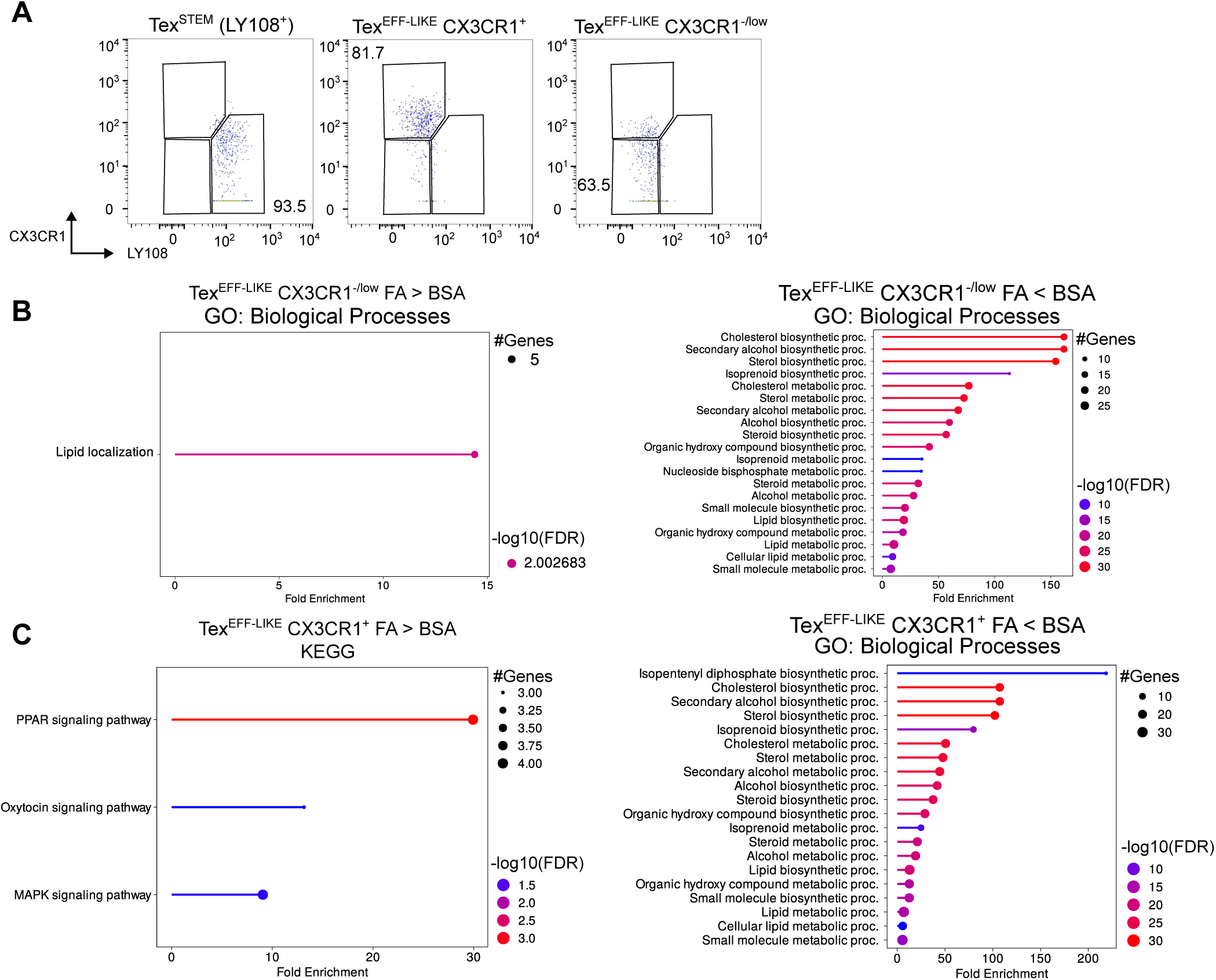
*Ex vivo* FA exposure alters lipid metabolism gene expression in Tex^INT^ and Tex^TERM^. C57BL/6 mice were infected with LCMV Cl13, and CD8^+^ PD-1^+^ Tex^STEM^ (LY108^+^ CX3CR1^-^) or Tex^EFF-LIKE^ (LY108^-^ CX3CR1^+^ or LY108^-^ CX3CR1^-/low^) splenocytes were FACS-purified at day 8 p.i.. (A) Flow cytometry showing CX3CR1 and LY108 expression in FACS-purified Tex subsets. (B-C) Tex subsets were incubated with palmitate-BSA (FA) or BSA alone (BSA) for 8 hours and processed for RNAseq. GO biological processes (B & C, right) and KEGG pathways (C, left) enriched by upregulated (left) or downregulated (right) genes in Tex^EFF-LIKE^ CX3CR1^-/low^ (B) and Tex^EFF-LIKE^ CX3CR1^+^ (C) after BSA vs. FA incubation. (A-C) Data pooled across 3 independent repeats with n=4-5 mice/experiment.

**Fig S4.**
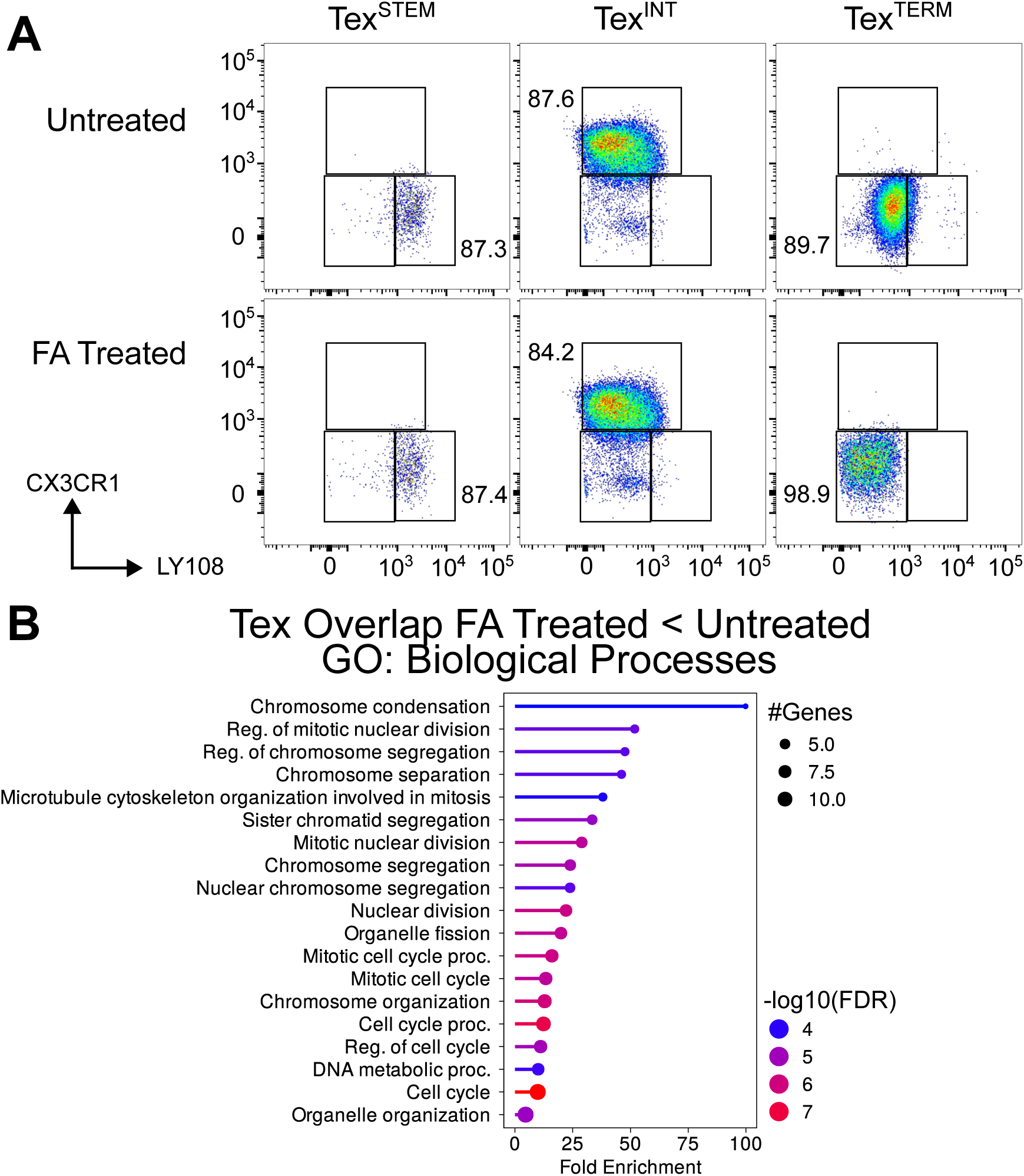
*In vivo* FA administration downregulated cell cycle related transcripts across Tex subsets from LCMV Cl13 infected mice. C57BL/6 mice were infected with LCMV Cl13 and treated i.p. with 0.3 mmol of FA (1:1 lauric and palmitic acid) or PBS every 12 hours from days 16 to 20 p.i.. CD8^+^ PD-1^+^ Tex^STEM^ (LY108^+^ CX3CR1^-^), Tex^INT^ (LY108^-^ CX3CR1^+^) or Tex^TERM^ (LY108^-^ CX3CR1^-^) splenocytes were FACS purified for RNAseq at day 21 p.i.. (A) Flow cytometry showing CX3CR1 and LY108 expression in FACS-purified Tex subsets. (B) GO analysis showing biological processes enriched by genes that were overlapping in all three Tex subsets and were downregulated in FA treated vs. untreated mice. Data combined across three experiments with n=4-5 mice/group.

**Fig S5.**
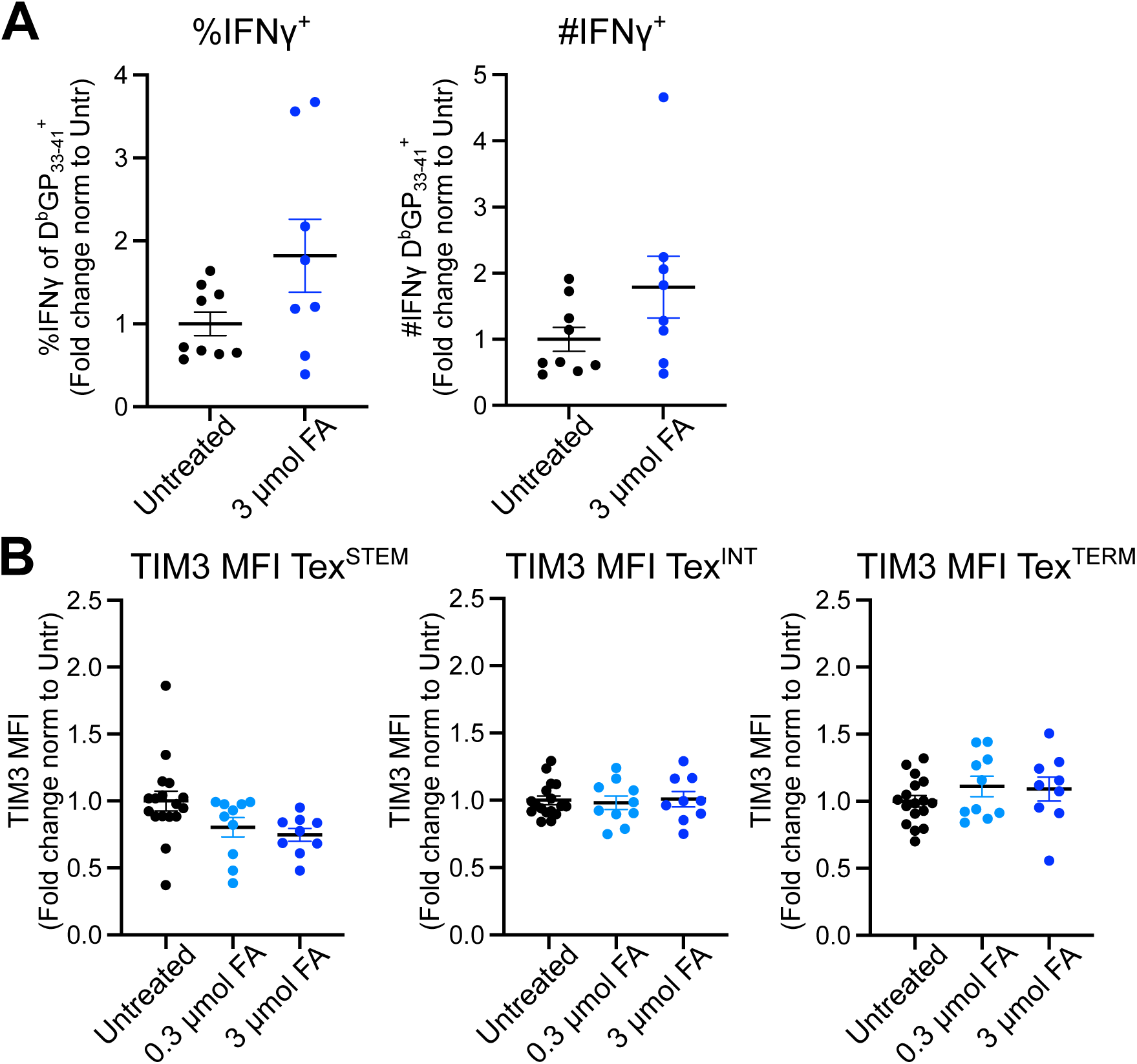
*In vivo* FA administration did not impact IFN-γ production or TIM3 expression during LCMV Cl13 infection. C57BL/6 mice were infected with LCMV Cl13 and treated i.p. with either 0.3 mmol or 3 mmol of FA (1:1 lauric and palmitic acid mix) or PBS every 12 hours from days 16 to 20 p.i.. Splenocytes were harvested and analyzed at day 21 p.i. (A) Numbers (left) and percentages (right) of interferon-γ producing CD8 T cells after 5 hr *ex vivo* stimulation with LCMV GP_33-41_ peptide. (B) Expression of TIM3 expression in CD8^+^ D^b^/GP_33-41_ tetramer^+^ Tex^STEM^ (LY108^+^ CX3CR1^-^) (left), Tex^INT^ (LY108^-^ CX3CR1^+^) (middle) or Tex^TERM^ (LY108^-^ CX3CR1^-^) (right) normalized to average from untreated mice. (A-B) Averages ± SEM. Data are combined across three experiments with n = 4-5 mice/group. (A) Unpaired two-tailed Student’s t-test. (B) One-way ANOVA with Tukey’s correction.

## References

1. J. M. Kulinski, V. L. Tarakanova, J. Verbsky, Regulation of antiviral CD8 T-cell responses. Crit Rev Immunol 33, 477–488 (2013).

2. H. W. Virgin, E. J. Wherry, R. Ahmed, Redefining chronic viral infection. Cell 138, 30–50 (2009).

3. L. M. McLane, M. S. Abdel-Hakeem, E. J. Wherry, CD8 T Cell Exhaustion During Chronic Viral Infection and Cancer. Annu Rev Immunol 37, 457–495 (2019).

4. M. Hashimoto et al., CD8 T Cell Exhaustion in Chronic Infection and Cancer: Opportunities for Interventions. Annu Rev Med 69, 301–318 (2018).

5. A. J. Zajac et al., Viral immune evasion due to persistence of activated T cells without effector function. J Exp Med 188, 2205–2213 (1998).

6. A. Gallimore et al., Induction and exhaustion of lymphocytic choriomeningitis virus-specific cytotoxic T lymphocytes visualized using soluble tetrameric major histocompatibility complex class I-peptide complexes. J Exp Med 187, 1383–1393 (1998).

7. E. J. Wherry, J. N. Blattman, K. Murali-Krishna, R. van der Most, R. Ahmed, Viral persistence alters CD8 T-cell immunodominance and tissue distribution and results in distinct stages of functional impairment. J Virol 77, 4911–4927 (2003).

8. H. Shin, E. J. Wherry, CD8 T cell dysfunction during chronic viral infection. Curr Opin Immunol 19, 408–415 (2007).

9. P. Klenerman, A. Hill, T cells and viral persistence: lessons from diverse infections. Nat Immunol 6, 873–879 (2005).

10. P. P. Lee et al., Characterization of circulating T cells specific for tumor-associated antigens in melanoma patients. Nat Med 5, 677–685 (1999).

11. Y. Iwai et al., Involvement of PD-L1 on tumor cells in the escape from host immune system and tumor immunotherapy by PD-L1 blockade. Proc Natl Acad Sci U S A 99, 12293–12297 (2002).

12. E. J. Wherry, D. L. Barber, S. M. Kaech, J. N. Blattman, R. Ahmed, Antigen-independent memory CD8 T cells do not develop during chronic viral infection. Proc Natl Acad Sci U S A 101, 16004–16009 (2004).

13. B. Bengsch et al., Bioenergetic Insufficiencies Due to Metabolic Alterations Regulated by the Inhibitory Receptor PD-1 Are an Early Driver of CD8(+) T Cell Exhaustion. Immunity 45, 358–373 (2016).

14. N. E. Scharping et al., The Tumor Microenvironment Represses T Cell Mitochondrial Biogenesis to Drive Intratumoral T Cell Metabolic Insufficiency and Dysfunction. Immunity 45, 374–388 (2016).

15. Y. R. Yu et al., Disturbed mitochondrial dynamics in CD8+ TILs reinforce T cell exhaustion. Nat Immunol 21, 1540–1551 (2020).

16. S. A. Vardhana et al., Impaired mitochondrial oxidative phosphorylation limits the self-renewal of T cells exposed to persistent antigen. Nat Immunol 21, 1022–1033 (2020).

17. D. T. Utzschneider et al., T Cell Factor 1-Expressing Memory-like CD8(+) T Cells Sustain the Immune Response to Chronic Viral Infections. Immunity 45, 415–427 (2016).

18. S. J. Im et al., Defining CD8+ T cells that provide the proliferative burst after PD-1 therapy. Nature 537, 417–421 (2016).

19. J. C. Beltra et al., Developmental Relationships of Four Exhausted CD8. Immunity 52, 825–841.e828 (2020).

20. E. Ahn, B. Youngblood, J. Lee, S. Sarkar, R. Ahmed, Demethylation of the PD-1 Promoter Is Imprinted during the Effector Phase of CD8 T Cell Exhaustion. J Virol 90, 8934–8946 (2016).

21. J. R. Giles et al., Shared and distinct biological circuits in effector, memory and exhausted CD8. Nat Immunol 23, 1600–1613 (2022).

22. M. S. Abdel-Hakeem et al., Epigenetic scarring of exhausted T cells hinders memory differentiation upon eliminating chronic antigenic stimulation. Nat Immunol 22, 1008–1019 (2021).

23. D. T. Utzschneider et al., Early precursor T cells establish and propagate T cell exhaustion in chronic infection. Nat Immunol 21, 1256–1266 (2020).

24. Z. Chen et al., TCF-1-Centered Transcriptional Network Drives an Effector versus Exhausted CD8 T Cell-Fate Decision. Immunity 51, 840–855.e845 (2019).

25. Y. A. Leong et al., CXCR5(+) follicular cytotoxic T cells control viral infection in B cell follicles. Nat Immunol 17, 1187–1196 (2016).

26. T. Wu >et al., The TCF1-Bcl6 axis counteracts type I interferon to repress exhaustion and maintain T cell stemness. Sci Immunol 1 (2016).

27. R. He et al., Follicular CXCR5-expressing CD8(+) T cells curtail chronic viral infection. Nature 537, 412–428 (2016).

28. W. H. Hudson et al., Proliferating Transitory T Cells with an Effector-like Transcriptional Signature Emerge from PD-1+ Stem-like CD8+ T Cells during Chronic Infection. Immunity 51, 1043–1058.e1044 (2019).

29. R. Zander et al., CD4 T Cell Help Is Required for the Formation of a Cytolytic CD8+ T Cell Subset that Protects against Chronic Infection and Cancer. Immunity 51, 1028–1042.e1024 (2019).

30. D. Zehn, R. Thimme, E. Lugli, G. P. de Almeida, A. Oxenius, ’Stem-like’ precursors are the fount to sustain persistent CD8. Nat Immunol 23, 836–847 (2022).

31. S. S. Gabriel et al., Transforming growth factor-β-regulated mTOR activity preserves cellular metabolism to maintain long-term T cell responses in chronic infection. Immunity 54, 1698–1714.e1695 (2021).

32. E. J. Wherry, M. Kurachi, Molecular and cellular insights into T cell exhaustion. Nat Rev Immunol 15, 486–499 (2015).

33. C. M. Metallo, M. G. Vander Heiden, Understanding metabolic regulation and its influence on cell physiology. Mol Cell 49, 388–398 (2013).

34. M. M. Rinschen, J. Ivanisevic, M. Giera, G. Siuzdak, Identification of bioactive metabolites using activity metabolomics. Nat Rev Mol Cell Biol 20, 353–367 (2019).

35. R. J. Cordy, et al., Distinct amino acid and lipid perturbations characterize acute versus chronic malaria. JCI Insight 4 (2019).

36. J. C. Schoeman et al., Metabolic characterization of the natural progression of chronic hepatitis B. Genome Med 8, 64 (2016).

37. A. L. Willig, E. T. Overton, Metabolic Complications and Glucose Metabolism in HIV Infection: A Review of the Evidence. Curr HIV/AIDS Rep 13, 289–296 (2016).

38. C. X. Zhou et al., Metabolomic Profiling of Mice Serum during Toxoplasmosis Progression Using Liquid Chromatography-Mass Spectrometry. Sci Rep 6, 19557 (2016).

39. A. Lercher et al., Type I Interferon Signaling Disrupts the Hepatic Urea Cycle and Alters Systemic Metabolism to Suppress T Cell Function. Immunity 51, 1074–1087.e1079 (2019).

40. L. K. Beura et al., Lymphocytic choriomeningitis virus persistence promotes effector-like memory differentiation and enhances mucosal T cell distribution. J Leukoc Biol 97, 217–225 (2015).

41. S. Li et al., Predicting network activity from high throughput metabolomics. PLoS Comput Biol 9, e1003123 (2013).

42. M. Wang et al., Sharing and community curation of mass spectrometry data with Global Natural Products Social Molecular Networking. Nat Biotechnol 34, 828–837 (2016).

43. L. W. Sumner et al., Proposed minimum reporting standards for chemical analysis Chemical Analysis Working Group (CAWG) Metabolomics Standards Initiative (MSI). Metabolomics 3, 211–221 (2007).

44. M. Wang et al., Mass spectrometry searches using MASST. Nat Biotechnol 38, 23–26 (2020).

45. H. Baazim et al., CD8+ T cells induce cachexia during chronic viral infection. Nat Immunol 20, 701–710 (2019).

46. L. Labarta-Bajo et al., CD8 T cells drive anorexia, dysbiosis, and blooms of a commensal with immunosuppressive potential after viral infection. Proc Natl Acad Sci U S A 117, 24998–25007 (2020).

47. F. Pietrocola et al., Metabolic effects of fasting on human and mouse blood in vivo. Autophagy 13, 567–578 (2017).

48. E. L. Pearce, M. C. Poffenberger, C. H. Chang, R. G. Jones, Fueling immunity: insights into metabolism and lymphocyte function. Science 342, 1242454 (2013).

49. G. J. van der Vusse, Albumin as fatty acid transporter. Drug Metab Pharmacokinet 24, 300–307 (2009).

50. C. Indiveri et al., The mitochondrial carnitine/acylcarnitine carrier: function, structure and physiopathology. Mol Aspects Med 32, 223–233 (2011).

51. S. M. Houten, S. Violante, F. V. Ventura, R. J. Wanders, The Biochemistry and Physiology of Mitochondrial Fatty Acid β-Oxidation and Its Genetic Disorders. Annu Rev Physiol 78, 23–44 (2016).

52. N. Longo, C. Amat di San Filippo, M. Pasquali, Disorders of carnitine transport and the carnitine cycle. Am J Med Genet C Semin Med Genet 142C, 77–85 (2006).

53. D. Montaigne, L. Butruille, B. Staels, PPAR control of metabolism and cardiovascular functions. Nat Rev Cardiol 18, 809–823 (2021).

54. M. Pasello, A. M. Giudice, K. Scotlandi, The ABC subfamily A transporters: Multifaceted players with incipient potentialities in cancer. Semin Cancer Biol 60, 57–71 (2020).

55. J. A. Menendez, R. Lupu, Fatty acid synthase and the lipogenic phenotype in cancer pathogenesis. Nat Rev Cancer 7, 763–777 (2007).

56. N. S. Chandel, Lipid Metabolism. Cold Spring Harb Perspect Biol 13 (2021).

57. M. Wu et al., Multiparameter metabolic analysis reveals a close link between attenuated mitochondrial bioenergetic function and enhanced glycolysis dependency in human tumor cells. Am J Physiol Cell Physiol 292, C125–136 (2007).

58. W. Pendergrass, N. Wolf, M. Poot, Efficacy of MitoTracker Green and CMXrosamine to measure changes in mitochondrial membrane potentials in living cells and tissues. Cytometry A 61, 162–169 (2004).

59. L. D. Zorova et al., Mitochondrial membrane potential. Anal Biochem 552, 50–59 (2018).

60. V. A. Boussiotis, Molecular and Biochemical Aspects of the PD-1 Checkpoint Pathway. N Engl J Med 375, 1767–1778 (2016).

61. M. Veldhoen, C. Ferreira, Influence of nutrient-derived metabolites on lymphocyte immunity. Nat Med 21, 709–718 (2015).

62. A. Lercher, H. Baazim, A. Bergthaler, Systemic Immunometabolism: Challenges and Opportunities. Immunity 53, 496–509 (2020).

63. H. Chi, Immunometabolism at the intersection of metabolic signaling, cell fate, and systems immunology. Cell Mol Immunol 19, 299–302 (2022).

64. W. Wasyluk, A. Zwolak, Metabolic Alterations in Sepsis. J Clin Med 10 (2021).

65. K. C. Fearon, D. J. Glass, D. C. Guttridge, Cancer cachexia: mediators, signaling, and metabolic pathways. Cell Metab 16, 153–166 (2012).

66. H. J. Patel, B. M. Patel, TNF-α and cancer cachexia: Molecular insights and clinical implications. Life Sci 170, 56–63 (2017).

67. S. K. Fried, R. Zechner, Cachectin/tumor necrosis factor decreases human adipose tissue lipoprotein lipase mRNA levels, synthesis, and activity. J Lipid Res 30, 1917–1923 (1989).

68. J. E. Rupert et al., Tumor-derived IL-6 and trans-signaling among tumor, fat, and muscle mediate pancreatic cancer cachexia. J Exp Med 218 (2021).

69. P. Matthys et al., Severe cachexia in mice inoculated with interferon-gamma-producing tumor cells. Int J Cancer 49, 77–82 (1991).

70. G. F. Grabner, H. Xie, M. Schweiger, R. Zechner, Lipolysis: cellular mechanisms for lipid mobilization from fat stores. Nat Metab 3, 1445–1465 (2021).

71. G. J. van der Windt et al., Mitochondrial respiratory capacity is a critical regulator of CD8+ T cell memory development. Immunity 36, 68–78 (2012).

72. G. J. van der Windt et al., CD8 memory T cells have a bioenergetic advantage that underlies their rapid recall ability. Proc Natl Acad Sci U S A 110, 14336–14341 (2013).

73. D. O’Sullivan et al., Memory CD8(+) T cells use cell-intrinsic lipolysis to support the metabolic programming necessary for development. Immunity 41, 75–88 (2014).

74. Y. Pan et al., Survival of tissue-resident memory T cells requires exogenous lipid uptake and metabolism. Nature 543, 252–256 (2017).

75. H. Frizzell et al., Organ-specific isoform selection of fatty acid-binding proteins in tissue-resident lymphocytes. Sci Immunol 5 (2020).

76. E. G. Hunt et al., Acetyl-CoA carboxylase obstructs CD8+ T cell lipid utilization in the tumor microenvironment. Cell Metab 36, 969–983.e910 (2024).

77. A. M. Prentice, Starvation in humans: evolutionary background and contemporary implications. Mech Ageing Dev 126, 976–981 (2005).

78. N. Collins et al., The Bone Marrow Protects and Optimizes Immunological Memory during Dietary Restriction. Cell 178, 1088–1101.e1015 (2019).

79. S. Xu et al., Uptake of oxidized lipids by the scavenger receptor CD36 promotes lipid peroxidation and dysfunction in CD8. Immunity 54, 1561–1577.e1567 (2021).

80. A. A. Al-Khami et al., Exogenous lipid uptake induces metabolic and functional reprogramming of tumor-associated myeloid-derived suppressor cells. Oncoimmunology 6, e1344804 (2017).

81. X. Ma et al., Cholesterol Induces CD8+ T Cell Exhaustion in the Tumor Microenvironment. Cell Metab 30, 143–156.e145 (2019).

82. Y. Zhang et al., Enhancing CD8+ T Cell Fatty Acid Catabolism within a Metabolically Challenging Tumor Microenvironment Increases the Efficacy of Melanoma Immunotherapy. Cancer Cell 32, 377–391.e379 (2017).

83. T. Manzo et al., Accumulation of long-chain fatty acids in the tumor microenvironment drives dysfunction in intrapancreatic CD8+ T cells. J Exp Med 217 (2020).

84. M. E. King et al., Long-chain polyunsaturated lipids associated with responsiveness to anti-PD-1 therapy are colocalized with immune infiltrates in the tumor microenvironment. J Biol Chem 299, 102902 (2023).

85. R. Ahmed, A. Salmi, L. D. Butler, J. M. Chiller, M. B. Oldstone, Selection of genetic variants of lymphocytic choriomeningitis virus in spleens of persistently infected mice. Role in suppression of cytotoxic T lymphocyte response and viral persistence. J Exp Med 160, 521–540 (1984).

86. C. A. Schneider, W. S. Rasband, K. W. Eliceiri, NIH Image to ImageJ: 25 years of image analysis. Nat Methods 9, 671–675 (2012).

87. M. C. Chambers et al., A cross-platform toolkit for mass spectrometry and proteomics. Nat Biotechnol 30, 918–920 (2012).

88. T. Pluskal, S. Castillo, A. Villar-Briones, M. Oresic, MZmine 2: modular framework for processing, visualizing, and analyzing mass spectrometry-based molecular profile data. BMC Bioinformatics 11, 395 (2010).

89. A. Gonzalez et al., Qiita: rapid, web-enabled microbiome meta-analysis. Nat Methods 15, 796–798 (2018).

90. J. A. Gilbert, J. K. Jansson, R. Knight, The Earth Microbiome project: successes and aspirations. BMC Biol 12, 69 (2014).

91. J. A. Vizcaíno et al., ProteomeXchange provides globally coordinated proteomics data submission and dissemination. Nat Biotechnol 32, 223–226 (2014).

92. J. Xia, D. S. Wishart, Web-based inference of biological patterns, functions and pathways from metabolomic data using MetaboAnalyst. Nat Protoc 6, 743–760 (2011).

93. R Core Team (2024) R: A Language and Environment for Statistical Computing. (R Foundation for Statistical Computing, Vienna, Austria).

94. R. C. Gentleman et al., Bioconductor: open software development for computational biology and bioinformatics. Genome Biol 5, R80 (2004).

95. M. I. Love, W. Huber, S. Anders, Moderated estimation of fold change and dispersion for RNA-seq data with DESeq2. Genome Biol 15, 550 (2014).

96. S. X. Ge, D. Jung, R. Yao, ShinyGO: a graphical gene-set enrichment tool for animals and plants. Bioinformatics 36, 2628–2629 (2020).

97. E. Ulgen, O. Ozisik, O. U. Sezerman, pathfindR: An R Package for Comprehensive Identification of Enriched Pathways in Omics Data Through Active Subnetworks. Front Genet 10, 858 (2019).

